# CAR modulates plasma membrane nano-organization and immune signaling downstream of RALF1-FERONIA signaling pathway

**DOI:** 10.1101/2021.10.13.464316

**Authors:** Weijun Chen, Huina Zhou, Fan Xu, Meng Yu, Alberto Coego, Lesia Rodriguez, Yuqing Lu, Qijun Xie, Qiong Fu, Jia Chen, Guoyun Xu, Dousheng Wu, Xiushan Li, Xiaojuan Li, Yvon Jaillais, Pedro L. Rodriguez, Sirui Zhu, Feng Yu

**Affiliations:** State Key Laboratory of Chemo/Biosensing and Chemometrics, Laboratory of Plant Functional Genomics and Developmental Regulation, College of Biology, Hunan University, Changsha 410082, P. R. Chin; Zhengzhou Tobacco Research Institute, Zhengzhou 450001, P.R. China; Beijing Advanced Innovation Center for Tree Breeding by Molecular Design, Beijing Forestry University, Beijing 100083, P. R. China; Instituto de Biología Molecular y Celular de Plantas, Consejo Superior de Investigaciones Científicas–Universidad Politécnica de Valencia, ES–46022 Valencia, Spain; Laboratoire Reproduction et Développement des Plantes, Université de Lyon, ENS de Lyon, CNRS, INRAE, F-69342 Lyon, France

**Keywords:** FER, C2 domain ABA–related proteins, lipid-binding, nanodomains, immune complex.

## Abstract

- The receptor-like kinase (RLK) FERONIA (FER) senses peptide ligands in the plasma membrane (PM), modulates plant growth and development, and integrates biotic and abiotic stress signaling for downstream adaptive response. However, the molecular interplay of these diverse processes is largely unknown.
- Here, we show that FER, the receptor of Rapid Alkalinization Factor 1 (RALF1), physically interacts with C2 domain ABA–related (CAR) proteins to control the nano-organization of the PM in *Arabidopsis*.
- During this process, the RALF1–FER pathway upregulates CAR protein translation, and then more CAR proteins are recruited to PM. This acts as a rapid feedforward loop to stabilize PM liquid-ordered phase. FER interacts with and phosphorylates CARs, and reducing their lipid-binding ability, which breaks the feedback regulation CAR activity at latter time point.
- Similar to FER mutant, pentuple *car14569* mutants inhibit flg22-induced FLS2-BAK1 immune complex formation, which depends on the integrity of nanodomains.
  - Together, we propose that the FER-CAR module controls the formation of the PM nano-organization during RALF signaling through a self-contained amplifying loop including both positive and negative feedbacks.

## Introduction

*Catharanthus roseus* receptor–like kinase 1–like protein (*Cr*RLK1L) FERONIA (FER) is a receptor-like kinase with versatile and tissue–specific functions in plant cell growth and survival. FER functions with its ligands, rapid alkalinization factors (RALFs, e.g., RALF1 and RALF23), as a regulator of fertilization (Escobar-Restrepo *et al*., 2007; Li *et al*., 2015), cell growth (Guo *et al*., 2009; Duan *et al*., 2010; Du *et al*., 2016; Li *et al*., 2018), hormone signaling (Guo *et al*., 2009; Duan *et al*., 2010), stress responses (Chen *et al*., 2016), immune signaling (Stegmann *et al*., 2017) and energy and RNA metabolism (Yang *et al*., 2015; Xu *et al*., 2019; Wang *et al*., 2020; Zhu *et al*., 2020). However, the mechanism through which FER quickly responds to and integrates external signals and transmits them to varied downstream targets remains unknown. Recent report showed that FER acts as a scaffold to regulate the formation of the immune receptor complex (FLS2 and BAK1) to respond to the pathogen–associated molecular pattern (PAMP) flagellin 22 (flg22) in a RALF–dependent manner (Stegmann *et al*., 2017), and this process depends on the regulation of RALF-FER on nanodomains (Gronnier *et al*., 2022).

Biological membranes are made of a plethora of co-existing nanodomains, which are characterized by their small size (i.e., below 1 µm) and concentration of related sterols and sphingolipids (Spira *et al*., 2012; Jarsch *et al*., 2014; Jaillais & Ott, 2020). Such membrane domains are critical in signaling notably because they can locally concentrate proteins, a process known as nanoclustering (Ott, 2017; Jaillais & Ott, 2020). By increasing the local concentration of proteins, nanoclustering favors the formation of protein complexes or post-translational modifications, which often act as a trigger to initiate signaling (Jaillais & Ott, 2020; Gronnier *et al*., 2022). The small protein family flotillin (Flot) and plant-specific remorin protein family have been described as marker proteins of nanodomains. Multiple evidences suggest that Flots and remorins are crucial in nanodomain stabilization and receptor recruitment into these structures, but the detailed molecular mechanism is still unknown (Liang *et al*., 2018; Huang *et al*., 2019; Jaillais & Ott, 2020; Pan *et al*., 2020). In addition, protein nanoclustering is regulated through the formation of various protein–lipid and protein– protein interactions (Simons & Gerl, 2010; Garcia-Parajo *et al*., 2014; Sezgin *et al*., 2017; Pan *et al*., 2020). As such, they act as versatile signaling platforms with very diverse composition in term of lipids and proteins (Burkart & Stahl, 2017; Platre *et al*., 2019; Jaillais & Ott, 2020). However, how plants regulate nanodomains by sensing the external environment signaling is just starting to be elucidated.

Accumulating evidence has shown that in plants, nanodomains are important for the cellular response to environmental cues and improving the fitness of plants (Demir *et al*., 2013; Wang *et al*., 2015; Bucherl *et al*., 2017; Huang *et al*., 2019). Several works have linked the abscisic acid (ABA) response with nanodomain dynamics (Demir *et al*., 2013; Diaz *et al*., 2016). For example, a 10-member family of lipid–binding C2 domain ABA–related proteins (CARs, CAR1 to CAR10) was identified and found to interact with pyrabactin resistance 1 (PYR1)/PYR1-like (PYL)/regulatory components of ABA receptors (RCAR) ABA receptors and to positively regulate ABA signaling (Rodriguez *et al*., 2014). CARs oligomerize at the PM in a calcium-dependent manner and has been identified located in nanodomains and induced membrane fusion via promoting membrane curvature (Martens & McMahon, 2008; Demir *et al*., 2013; Diaz *et al*., 2016). However, the upstream regulator(s) that link CARs with the modulation of the PM landscape is still unknown. Herein, we propose a working model in which FER works together with lipid-binding CAR proteins to regulate the partitioning of liquid-ordered and liquid-disordered phases. In this scenario, RALF1 signaling modulated the formation of higher density nanodomains to quickly respond to external signals by activating FER and regulating the accumulation and phosphorylation of downstream CAR proteins.

## Materials and Methods

### Plant Material and Growth Conditions

*Arabidopsis* (Col-0) was used as the wild-type (WT). The following transgenic and mutant lines were used in this study: *CAR1-Myc, CAR4-Myc, CAR5-Myc, CAR6-Myc, CAR9-Myc, CAR10-Myc,* CAR9-Myc/*fer-4, YFP–CAR9*, *YFP–CAR9*/*fer-4,* CAR9^3M^-YFP/*fer-4* and *car59, car145, car459, car149, car159, car1459, car14569–Δ1, car14569–Δ4, car14569–Δ10, car14569–Δ20*.

*Arabidopsis thaliana* seeds were first surface sterilized by 75 % (v/v) ethanol. After stigmatization at 4 °C for 3 d, seeds were growing on 1/2 MS with 0.8 % sucrose and 1.0 % Phytagel (Sigma–Aldrich) at 22 °C in a 16–h light/8–h dark condition for subsequent analysis. Details of this information are described in Supporting information methods S1.

### Yeast Two–Hybrid (Y2H) Assay

Y2H assays were performed as previously described (Du *et al*., 2016). Briefly, the cytoplasm domain of *Cr*RLK1L family members (AT3G51550, FER, 469–896 amino acids; AT2G39360, CVY1, 429–815 amino acids; AT5G38990, MDS1, 462–880 amino acids; AT5G39020, MDS3, 459–813 amino acids; AT1G30570, HERK2, 451– 849 amino acids), and two RLKs (AT5G46330, FLS2, 870–1155 amino acids; AT4G39400, BRI1, 814–1196 amino acids) were fused in–frame with the GAL4 DNA–binding domain of the bait vector pGBKT7. The CDS of *CARs* were fused with the GAL4 DNA–activating domain of the prey vector pGADT7 (AD–CAR). Distinct plasmid pairs were transformed into yeast AH109 cells. The diluted transformants were plated onto synthetic dropout medium lacking tryptophan/leucine (SD/–Trp–Leu+His) and synthetic dropout medium lacking tryptophan/leucine/histidine (SD/–Trp–Leu–His) but supplemented with 20 mM 3–amino–1,2,4–triazole (3–AT) for 4 days to test the interaction.

### Co–Immunoprecipitation (Co–IP) Assay

Co–IP was performed as previously described with some modifications (Li *et al*., 2018). Seven–day–old seedlings *35S::CARs–Myc* (about 0.5 g) were ground to a fine powder in liquid nitrogen and solubilized with 500 µL TBST buffer [50 mM Tris–HCl (pH 7.5), 150 mM NaCl, 5 mM MgCl_2_, 1 mM EDTA, 1 % Triton X–100] containing 1 × protease inhibitor cocktail (78430, Thermo Fisher Scientific) and 1 × phosphatase inhibitor (78420, Thermo Fisher Scientific) and spun for 1 hour at 4 °C. The extracts were centrifuged at 16,000 g at 4 °C for 10 min, and 500 µL supernatant was transferred to incubate with prepared 20 µL Anti–Myc magnetic beads (B26301) overnight at 4 °C, and 100 µL supernatant was used as input. Then, the beads were washed 3 times with the washing buffer [50 mM Tris–HCl (pH 7.5), 150 mM NaCl, 0.1 % Triton X–100] containing 1 × protease inhibitor cocktail and eluted with elution buffer (0.2 M Glycine, 1 % Triton X–100, 1 mM EDTA, pH 2.5). And proteins were run on a 10 % SDS– PAGE gel and analyzed by anti–FER (Du *et al*., 2016; Li *et al*., 2018) and anti–Myc (CST, 2276S) antibody.

### *In vitro* Phosphorylation Assay

The phosphorylation and dephosphorylation assays were described as previously (Du, *et al*., 2016). GST–CAR9, GST–CAR5, FER–CD–His and FER^K565R^–CD–His (kinase dead form) were purified for phosphorylation assay *in vitro*.

The ABA–induced phosphorylation co–expression system was described previously (Li *et al*., 2018). Vectors of pACYC–PYL1–FER (S–tag), pACYC–PYL1– FER^K565R^ (S–tag), and pRSF–ABI1–CAR9 (His–tag) were constructed and co– transformed into one BL21 (DE3) *E. coli* strain. His–CAR proteins were purified and then subjected to liquid chromatography–tandem mass spectrometry (LC–MS/MS). Details of this method are described in Supporting information methods S1.

### *In vivo* Phosphorylation Assay

Phos–tag assay was performed as described previously (Chen *et al*., 2018). Seven–day– old *CAR–Myc* seedlings were used to analyze phosphorylated band *in vivo*. For phosphorylation sites detecting *in vivo*, *CAR–Myc* was enriched by immunoprecipitate. The CAR–Myc bands digested and analyzed by LC–MS/MS as described previously. Details of this method are described in Supporting information methods S1.

### Pan–anti–CARs Antibody Production

Antibody production was performed as previous study (Li *et al*., 2018). This antibody is raised by using full–length CAR9 expressed and purified from *E.coli* BL21. A 1–month–old ICR mouse (SLAC laboratory animal) was injected with 50 μg GST–CAR9 protein emulsified with Complete Freund’s adjuvant (F5881, Sigma–Aldrich). Two weeks later, 50 µg GST–CAR9 protein emulsified with Incomplete Freund’s adjuvant (F5506, Sigma–Aldrich) was injected into the ICR mouse and then once again in the next week. The serum of the immunized mouse was obtained as antibody for immunoblot detection. We tested the antibody in WT, *car9*, *car459* and *car14569* mutant lines by an immunoblot. Because of ten CAR family members share common motif and sequence, we still detected homologs in CAR9 signal mutants using this antibody, but this interacted band will be sharply reduced when using in CARs multiple mutants. Thus, we named this antibody as pan–anti–CARs antibody.

### Lipid Binding Assay

Phospholipid binding to proteins was assessed as described (Rodriguez *et al*., 2014). Briefly, phosphatidylserine (PS): phosphatidylcholine (PC) (25:75) was prepared in chloroform and dried to obtain a thin layer under a stream of nitrogen. The dried lipids were resuspended in buffer A (100 mM NaCl, 50 mM HEPES, pH 6.8) and mixed by vortexing for 20 min. Large multilamellar vesicles were obtained by centrifugation at 16,000 g for 20 min. Resuspended the vesicles in 1 mL of buffer A with multiple concentrations of free Ca^2+^ and stored at 4 °C. The vesicles (about 100 μg of phospholipids) were mixed with the different forms of GST–CAR9 (5 μg). The mixture was incubated with or without 1 mM Ca^2+^ via gentle shaking (250 rpm) for 30 min at room temperature. The vesicles and the bound proteins were pelleted by centrifugation at 12,000 g at 4 °C for 10 min and washed twice with 0.5 mL of buffer A. Samples were boiled with 1 × protein loading buffer for 10 min and separated by 10 % SDS–PAGE. Proteins were revealed by using anti–GST antibody.

### Variable–Angle Total Internal Reflection Fluorescence Microscopy (VA–TIRFM) Imaging

VA–TIRFM was performed as previously described (Li et al., 2012). For observation, seedlings were softly transferred onto a glass slide and soaked with liquid 1/2 MS, then covered with a coverslip. A custom–built VA–TIRFM based on an Olympus IX–71 microscope equipped with a total internal reflective fluorescence illuminator and a 100 × oil–immersion objective (numerical aperture of 1.45, Olympus) was used here. The 473 nm laser line from a diode laser was used to excite GFP. The emission fluorescence signals were detected with a back–illuminated EM–CCD camera (ANDOR iXon DV8897D–CS0–VP; And / or Technology, Belfast, UK) after filtering with band–pass filters. The gain of the EMCCD camera was set to 300 throughout all single–molecule imaging experiments; the setting was in the linear dynamic range of the EMCCD camera. Images were acquired with an exposure time and analyzed with Image J.

### Polysome Profiling Assay and Real–Time PCR

*Arabidopsis* polysomes profiling assay was performed as described (Missra & Von Arnim, 2014; Zhu *et al*., 2020). Briefly, 7–day–old seedlings (about 0.5 g) were treated with RALF1 (1 μM) for 10 min and then ground in liquid nitrogen followed by resuspension in polysome extraction buffer. Supernatant was loaded onto a 15 %–60 % sucrose gradient and spun in a Beckman SW 32 Ti rotor at 170,000 g for 4 h at 4 °C. We manually collected 11 fractions. Then, isolated the RNA from the last four fractions by RNAiso plus (Takara) kit. RT–qPCR analysis was used to quantify the total target relative transcript content in the input and the relative content of target transcripts associated with heavier polysomes in the bottom four fractions. Primers for RT–qPCR is shown in Table S1. *ACTIN* as a reference gene to detect the RNA levels of CARs.

### Di–4–ANEPPDHQ Lipid Staining

For di–4–ANEPPDHQ lipid staining, four–day–old seedlings are collected for each genotype. 5 μM (work concentration) di–4–ANEPPDHQ was added in liquid 1/2 MS medium for staining. Quantification of lipid polarity in root cell has been described previously (Huang et al., 2019; Owen et al., 2011). Details of this method are described in Supporting information methods S1.

### ROS Burst Assay

The flg22*–*induced ROS burst measurement assay was performed as previously described (Stegmann *et al*., 2017). Four–week–old *Arabidopsis* rosette leaves were used for ROS dynamic determination. ROS fluorescent probe H2DCF-DA staining (James *et al*., 2015) 6-day-old seedlings were treated with the liquid 1/2 MS medium containing 10 μM H2DCF-DA. Fluorescence was monitored with a confocal microscope using an excitation wavelength of 488 nm. Details of this method are described in Supporting information methods S1.

### *Pto* DC3000 infection assay

For *Pto* DC3000 infection assay (Stegmann *et al*., 2017), *Pseudomonas syringae pv. Tomato (Pto)* DC3000 strains were grown overnight in King’s B medium (10 g/L proteose peptone, 1.5 g/L anhydrous K_2_HPO_4_, 5 g/L MgSO_4_) with shaking at 28°C. Bacteria were collected from centrifuge tube and resuspended in water containing 0.02% Silwet L77 (Sigma Aldrich) to an OD_600_ = 0.2 (10^8^ colony forming units per mL). This bacterial suspension was sprayed on 4-week-old plants, which were covered with vented lids for 3 days. Three leaf discs per sample from different plants were collected in different microfuge tubes and ground with a drill-adapted pestle. Serial dilutions were plated on LB agar and colonies were counted 2 days later.

### Statistical Analysis

Software SPSS Statistics 17.0 was used. Data are shown as mean ± s.d.; * *p* < 0.05, ** *p* < 0.01, *** *p* < 0.001; n.s., not significant. One–way ANOVA with Tukey’s test were used as mention in figure note.

## Results

### Lipid-binding CAR proteins physically interact with the receptor kinase FER at the PM

FER is involved in response to multiple extracellular signals (Zhu *et al*., 2021), and has been evidenced for regulation of proteins in nanodomains to respond to RALF signal (Gronnier *et al*., 2022), but how this works mechanistically is still unclear. To understand the functions of FER in regulating membrane lipids, we performed a yeast two-hybrid (Y2H) screen using the cytoplasmic domain of FER (FER–CD, 469–896 amino acids, aa) as bait (Du *et al*., 2016). Among the interacting proteins, we found *CAR9*, which belongs to a 10-member family of lipid–binding C2 domain ABA–related proteins (Rodriguez *et al*., 2014; Diaz *et al*., 2016; Qin *et al*., 2019). We cloned the full-length genes for the ten *CAR* family members (*CAR1 – CAR10*) and found that *CAR1, CAR4, CAR5, CAR6, CAR9* and *CAR10* were able to directly interact with FER–CD in Y2H assays (Fig. 1a and Fig. S1a). Next, glutathione S–transferase (GST) pull–down assays further confirmed the interaction between CARs (GST-tagged CAR5, CAR6, CAR9 and CAR10) and His-tagged FER–CD (Fig. 1b). Furthermore, bimolecular fluorescence complementation (BiFC) assays were used to test the interaction between CARs and FER in *Arabidopsis* protoplasts. The results showed that CAR1, CAR4, CAR5, CAR6, CAR9, and CAR10 interacted with FER at the PM (the protein–protein interaction caused green fluorescence, which overlapped with FM4–64 PM red fluorescence) (Fig. 1c and Fig. S1b). We next performed coimmunoprecipitation (co– IP) assays and confirmed the interaction between Myc–tagged CAR proteins (i.e., CAR1, CAR4, CAR5, CAR6, CAR9, CAR10) and FER *in vivo* (Fig. 1d and Fig. S1c).

**Fig. 1.**
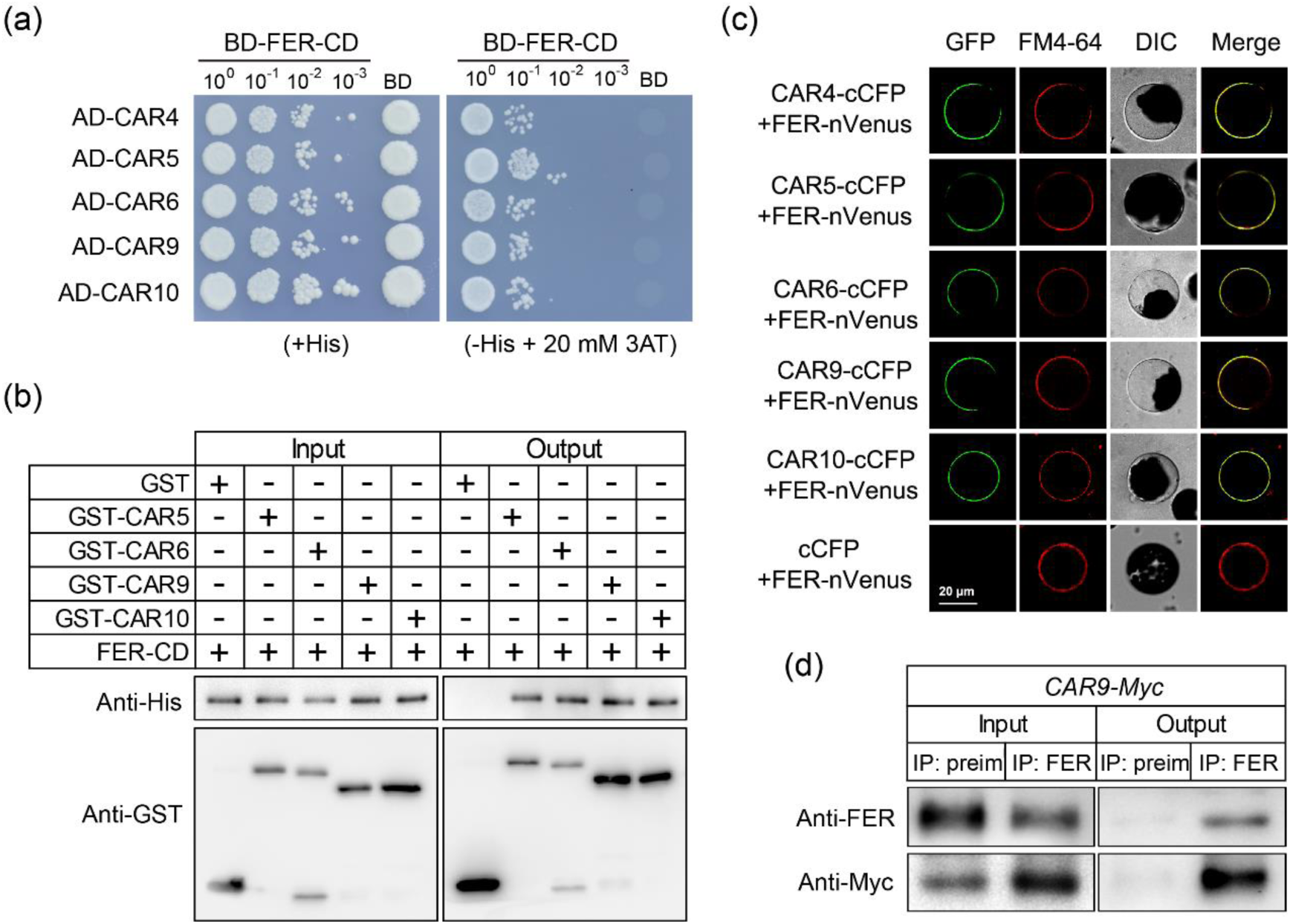
FER physically interacts with CARs. (a) Y2H assays showing the interaction between CARs and FER. SD (–His/–Leu/–Trp) selection medium containing 20 mM 3–AT was used to test the interaction. The CAR members were cloned into the pGADT7 (AD) vector, and the FER-CD was cloned into the pGBKT7 (BD) vector. (b) GST pull–down assay. The indicated GST–tag and His–tag were detected by anti– GST and anti–His, respectively. (c) FER interacts with CARs in BiFC assays in Arabidopsis protoplasts. Negative control (cCFP + FER–Venus) are shown, and FM4–64 indicates the PM (red). (d) Co–IP assays. The immunoprecipitated CAR9 and coimmunoprecipitated FER were indicated using anti–FER and anti–Myc antibodies, respectively. At least three biological replicates of (a) – (d) were performed with similar results.

To further verify the specificity of the interaction between CAR and FER, we cloned the cytoplasmic domains of five *Cr*RLK1L family members (CVY1, AT2G39360; BUPS1, AT4G39110; MDS1, AT5G38990; MDS3, AT5G39020; HERK2, AT1G30570) and two LRR-RLKs (FLS2 and BRI1) and further analyzed their interaction with CAR9. The results of the Y2H assays showed that CAR9 interacted specifically with FER and BUPS1, but not with the other six kinase domains (Fig. S1d). We also performed co–IP assays between Y2H noninteracting CARs (e.g., CAR3 and CAR7) and FER (Fig. S1e). The results revealed no interaction of CAR3 and CAR7 with FER *in vivo*, which indicates that the interaction between certain CARs and FER is specific. Taken together, these data suggest that FER physically interacts with a subset of CAR proteins at the PM.

### FER phosphorylates CARs, which further regulates the interaction of CARs with lipids

To evaluate whether CARs could be substrates of FER, we first tested CAR5 and CAR9 by using phosphorylation assays *in vitro*. The results showed that recombinant FER– CD induced a higher phosphorylation level in the CAR5 and CAR9 proteins (Fig. 2a). To further assess whether the phosphorylation levels of these two proteins were enhanced by RALF1 *in vivo*, Phos–Tag–PAGE analysis (Chen *et al*., 2018) was used to detect phosphorylated forms of CAR5 and CAR9 (there after referred to as pCAR5 and pCAR9) in plants expressing Myc-tagged CARs with or without RALF1 treatment (Fig. 2b). Notably, RALF1 treatment altered CAR protein abundance in *Arabidopsis* (detailed description in Fig. 3); thus, we adjusted the total protein levels (including those of phosphorylated and dephosphorylated proteins). We found a slow running form of CAR5 and CAR9, that was strongly reduced by alkaline phosphatase treatment (CIP), suggesting that they correspond to pCAR5 and pCAR9. RALF1 treatment (1 µM, 30 min) increased the phosphorylation levels of both CAR5 and CAR9 *in vivo* (Fig. 2b). Time series analysis indicated that the phosphorylation level of CAR5 and CAR9 increased at 20 min and 40 min post RALF1 treatment (protein abundance was adjusted prior to western blotting) in the WT background, but not in *fer-4* (Fig. S2). Together, our results indicate that RALF1-mediated CAR phosphorylation is dependent on FER. We further identified the FER-mediated phosphorylation sites of CAR5 and CAR9 using a previously developed ABA-induced co-expression system (Li *et al*., 2018). A kinase-dead mutant of FER–CD (Lys at 565 was mutated into Arg, FER–CD^K565R^) was used as a negative control (Escobar-Restrepo *et al*., 2007). Electrospray ionization (ESI) mass spectrometry (ESI-MS) analysis indicated that after ABA induction, FER phosphorylated CAR5 at Ser61, Ser62, and Tyr65 (Fig. S3a–S3c). Of note, in the same assay, Thr26, Ser27, and Tyr30 were phosphorylated in CAR9 (Fig. S3d and S3e), which are the equivalent phosphorylated residues in CAR5 (Fig. S4), reinforcing the idea that those phosphorylation sites could be physiologically relevant. Moreover, 2 out of 3 sites were further verified *in vivo* by label–free quantitative MS, and we found that RALF1 enhanced the CAR5 phosphorylation level at Ser62 and Tyr65 compared with mock-treated plants (Fig. 2c and Fig. S3f). Next, we mutated the three amino acid residues of CAR5 and CAR9 to either alanine (Ala) or phenylalanine (Phe) (GST– CAR5^3M^/9^3M^, Ser and Thr were mutated into Ala, Tyr was mutated into Phe) (Li *et al*., 2018; Zhou *et al*., 2018) to test whether GST–CAR5^3M^ and GST–CAR9^3M^ were phosphorylated by FER–CD. As expected, GST–CAR5^3M^ and GST–CAR9^3M^ remained unphosphorylated by FER (Fig. 2d). These data suggest that RALF1 promotes the FER-mediated phosphorylation of CAR5 and CAR9 at specific sites.

**Fig. 2.**
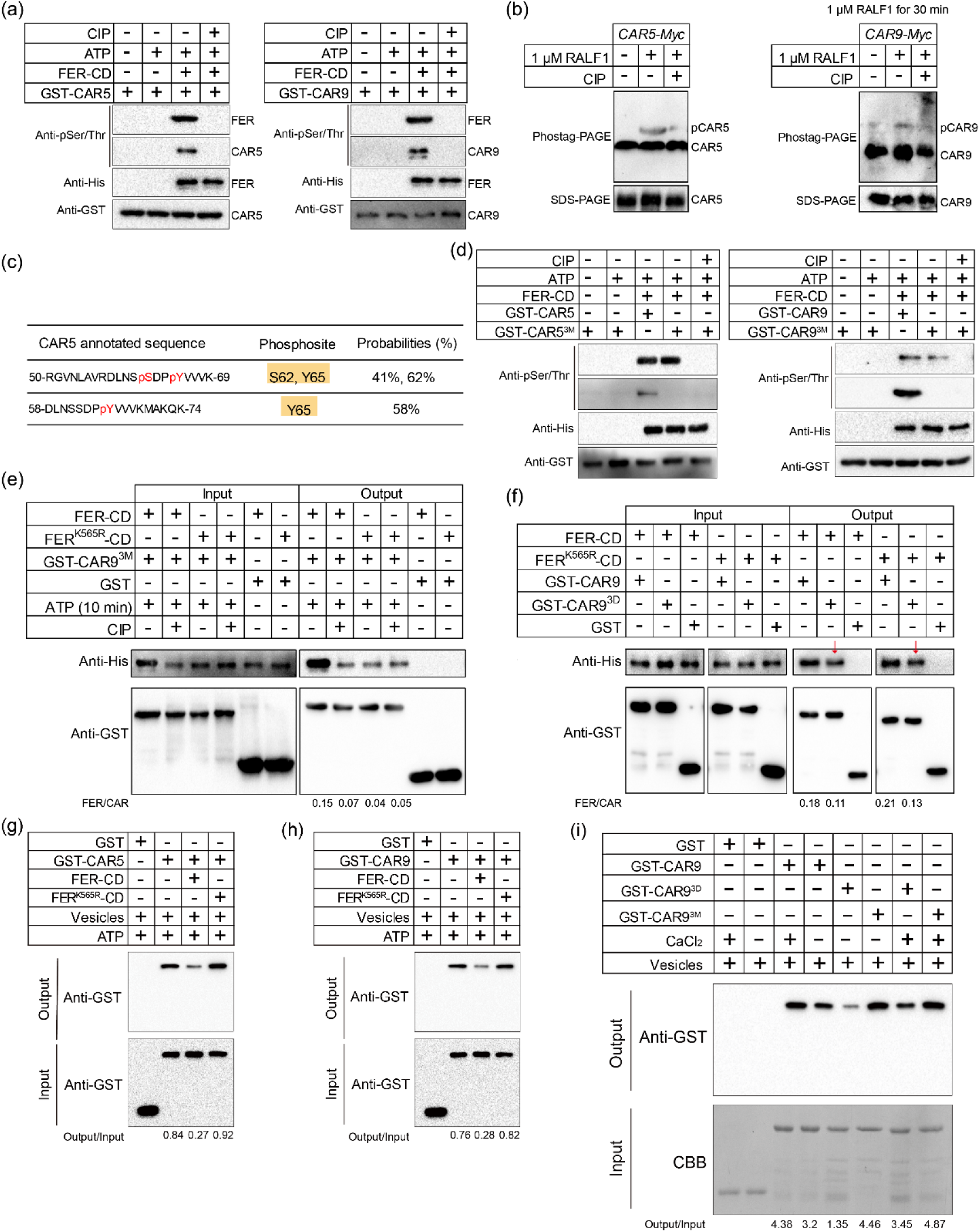
RALF1–FER phosphorylates CAR proteins and further regulates the interaction of CAR proteins and lipid. (a) *In vitro* phosphorylation assays showing that FER–CD can phosphorylate GST– CAR5 and GST–CAR9, as detected phosphorylation by using anti–pSer/Thr, and anti– GST and anti–His was used to detect CAR and FER-CD proteins. (b) Phostag–PAGE assays *in vivo*. Phosphorylated CAR5–Myc and CAR9–Myc bands are indicated as pCAR5–Myc and pCAR9–Myc. (c) Examples of phosphopeptides of CAR5 *in vivo*. The maximum probability shown for each phosphorylation site was calculated by the Andromeda algorithm integrated in MaxQuant. (d) *In vitro* phosphorylation assays showing that GST–CAR5^3M^ and GST–CAR9^3M^ were not phosphorylated by FER–CD. (e) GST pull–down assay showing that phosphorylated FER–CD exhibits a higher affinity towards GST–CAR9^3M^. The relative protein intensity of FER–CD–His and FER^K565R^–CD–His in the output was analyzed using ImageJ. (f) GST pull–down assay. GST–CAR9^3D^ shows a weaker interaction with FER–CD– His and FER^K565R^–CD–His. The relative protein intensity of FER–CD–His and FER^K565R^–CD–His in the output was analyzed using ImageJ. The red arrow pointing the weakened anti-His panel. (g-h) Phosphorylated CAR5 and CAR9 (0.1 μM) show weaker lipid binding ability in the presence of 1 mM CaCl_2_. (i) Lipid binding abilities of three forms of CAR9 in the presence or absence of 1 mM CaCl_2_. Immunoblot assays showing the protein levels by ImageJ. At least three biological replicates of (a) – (i) were performed with similar results.

**Fig. 3.**
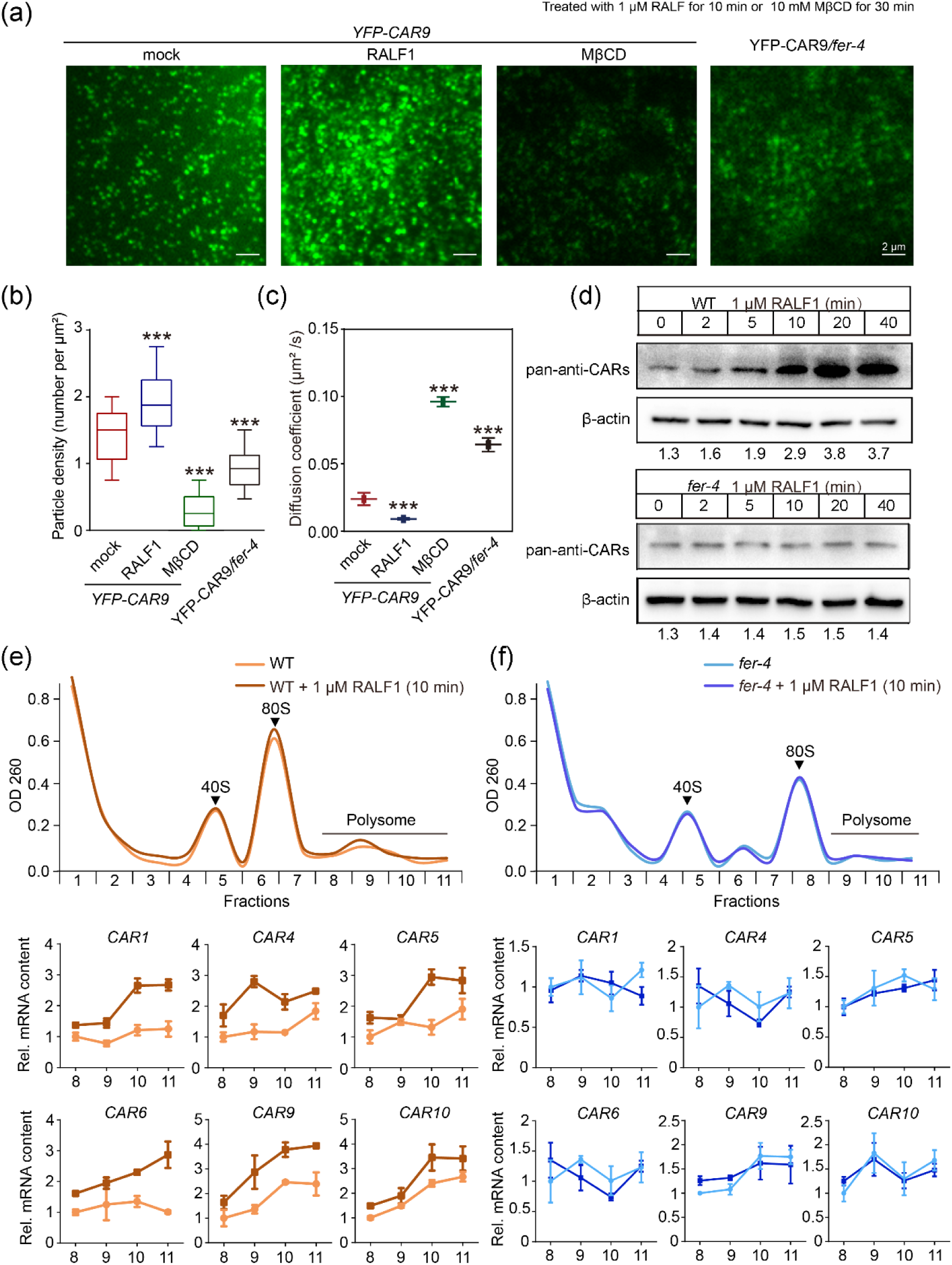
RALF1-FER promotes the formation of CAR-nanoclustering by enhancing their translation rate. (a) VA-TIRFM images of YFP-CAR9 at the PM in root cells with or without RALF1 (1 μM, 10 min) or MβCD (10 mM, 30 min) treatment in WT and *fer-4*background. (b) Quantitative comparison of YFP-CAR9 particle density in different groups. (c)) Comparison of diffusion coefficients of YFP-CAR9 particles in different groups. (d) RALF1–induced accumulation of CAR proteins in WT and *fer-4*. CAR proteins were detected by pan–anti–CARs antibody (see Methods for details of this antibody). The relative accumulation of CAR proteins was analyzed using ImageJ. (e-f) Upper panel: ribosome profiles. Fractions (8–11) containing mRNAs associated with polysomes are indicated with a black line. Lower panels: RT–qPCR results of polysome–associated CAR mRNAs. At least three biological replicates were performed with similar results. Data are shown as the mean ± s.d., ***p < 0.001, n.s., not significant. One–way ANOVA with Tukey’s test.

Further analysis of the phosphorylation levels of FER and CAR proteins on their function, we found that FER–CD (+ ATP) (ATP activates FER self–phosphorylation) showed a higher affinity toward CAR9^3M^ than the kinase–dead FER–CD^K565R^ (+ ATP) form (Fig. 2e). In addition, CIP reduced the interaction of FER–CD (+ ATP) but not FER–CD^K565R^ (+ ATP) with CAR9^3M^ (Fig. 2e). We then tested the effect of a CAR9 phosphorylation mimic on this interaction. We mutated three phosphorylation sites of CAR9 (Thr26, Ser27, Tyr30) to Asp (D) (GST–CAR9^3D^ mimics phosphorylated CAR9) to test how the mutant residues affected the FER–CAR9 association. Compared with wild-type CAR9, GST–CAR9^3D^ exhibited a lower association with FER–CD (41% ± 5.2% weakened) and FER^K565R^–CD (42% ± 7.4% weakened) (Fig. 2f). These results indicated that the phosphorylation of FER promotes the FER–CAR9 interaction, whereas the phosphorylation of CAR9 reduces its affinity toward FER.

CARs have been found located in nanodomains and directly interact with negatively charged phospholipid to promote membrane fusion (Martens *et al*., 2007; Demir *et al*., 2013; Rodriguez *et al*., 2014). To test whether FER regulates the interaction between CAR and lipids via phosphorylation, we investigated the lipid– binding ability of CAR under phosphorylated and nonphosphorylated states (Rodriguez *et al*., 2014). Since the Ca ^2+^-dependent anionic phospholipid binding is a hallmark of many C2 domains (Rodriguez *et al*., 2014), we confirmed that several CAR proteins (CAR5, 6, 9, and 10) directly interact with negatively charged phospholipid vesicles [25:75 (w/w) mixture of phosphatidylserine: phosphatidylcholine] in the presence of free Ca^2+^ (Fig. S5a). Furthermore, we compared the effect of wild-type FER and kinase-dead FER (FER^K565R^) on CAR5 and CAR9 in lipid cosedimentation assays and found that phosphorylated CAR5 and CAR9 (coincubated with FER–CD and ATP) had a weakened capacity to interact with anionic phospholipid (Fig. 2g and 2h). Furthermore, comparison of the lipid binding ability of two mutant forms of CAR9 protein (GST– CAR9^3M^ and GST–CAR9^3D^) toward negatively charged phospholipid vesicles revealed that CAR9^3M^ exhibited a similar association with vesicles than the wild-type CAR9, whereas CAR9^3D^ showed weaker binding ability (Fig. 2i).

A previous study used negative stain transmission electron microscopy (TEM) to show that CAR can interact with negatively charged phospholipid vesicles and further induce membrane curvature (Diaz *et al*., 2016). Evidence for membrane curvature enables lipid anchor proteins sorting to liquid-ordered membrane phases has recently emerged (Larsen *et al*., 2015). To answer whether FER could modulate membrane curvature via regulating the affinity between CARs and anionic phospholipid, we tested the effect of phosphorylation activity on CAR9-induced membrane curvature (Fig. S5b–S5i). Tubulation of liposomes has been widely used to study membrane curvature *in vitro* (Martens *et al*., 2007; Diaz *et al*., 2016). Addition of free Ca^2+^ and CAR9 to negatively charged phospholipid vesicles induced strong tubulation of the membranes (Fig. S5f and S5i). To investigate whether the phosphorylation could impact CAR9 to induce membrane curvature, the coincubation of GST–CAR9^3M^ or GST–CAR9^3D^ with vesicles was tested. From the results, GST–CAR9^3M^ induced strong tubulation of liposomes (Fig. S5g and S5i), whereas GST–CAR9^3D^ significantly reduced tubulation (Fig. S5h and S5i), which may suggest that RALF1-FER participates in the regulation of membrane curvature by regulating the phosphorylation of CAR protein. Taken together, FER phosphorylates CARs at specific sites, which further fine-tunes the association of FER and CARs. Moreover, as a lipid binding protein, CAR9-induced membrane curvature is finely regulated by the kinase activity of FER.

### RALF1-FER regulates CAR nanoclustering via promoting CAR protein translation

From above, several evidences implied that CAR proteins located in nanodomains are regulated by FER. To investigate how RALF1-FER affects the CAR nanoclusters on the PM, time-lapse images were collected with a variable–angle total internal reflection fluorescence (VA-TIRF) microscope for 4-day-old *YFP-CAR9* root cells followed by single-particle-tracking method, which is commonly used to investigate protein nanocluster (Pan *et al*., 2020). Treatment with 1 μM RALF1 resulted in a dramatic increase in the density of YFP-CAR9 particles (Fig. 3a and Fig. 3b), which suggests that more CAR proteins are localized to the membrane and recruited into CAR-associated nanoclusters. Furthermore, similar to the treatment with sterol-depleting agent methyl–β–cyclodextrin (MβCD), density of YFP-CAR9 particles in *fer-4* background was showed significantly decreased (Fig. 3a and Fig. 3b). Evidence shows that many proteins on PM are not uniformly mixed with membrane lipids via free diffusion-based equilibrium, but laterally segregated into nanodomains with a limited diffusion and specific lipids (Jaillais & Ott, 2020; Pan *et al*., 2020). Thus, we tested diffusion rate and revealed that treatment with RALF1 induced a slower diffusion coefficient of YFP-CAR9 (Fig. 3c), and on the contrary, YFP-CAR9 showed a highly diffusion coefficient after treatment with MβCD, in line with YFP-CAR9 under *fer-4* background (Fig. 3c). Moreover, we assessed the impact of the phosphorylation level of CAR9 on its nanoclustering. We expressed the YFP-CAR9^3D^ and YFP-CAR9^3M^ in WT background. VA-TIRFM analysis showed that YFP-CAR9^3D^ particles resulted in a shorter dwell time than YFP-CAR9 and YFP-CAR9^3M^ (Fig. S6). These results suggest that RALF1 promotes the formation of CAR9 nanoclusters, which possibly by recruiting more CAR proteins into nanodomains and coalescing small nanoclusters. Subsequently, FER negatively regulates the dwell time of CAR proteins in nanodomain via directly interacting with and phosphorylating CAR on the membrane, thus forming a feedback loop.

Ca^2+^-dependent C2 domains allow to be recruited to PM via interacting with negatively charged phospholipids (Martens *et al*., 2007). Thus, RALF1 promotes the formation of CAR9 nanoclusters, which means that more CAR9 proteins in the cytoplasm are recruited into PM. A recent study showed that the RALF1–FER promotes protein synthesis through facilitating translation initiation (Zhu *et al*., 2020); thus, we hypothesis that RALF1–FER facilitates the formation of CAR-related nanoclusters via promoting their protein synthesis. To test this hypothesis, we investigated the protein accumulation of CARs by using a pan–anti–CARs antibody (see Methods for details) (Fig. S7a) and found that RALF1 triggered the rapid accumulation of CAR proteins in WT, whereas the protein accumulation of CARs in the *fer–4* mutant was insensitive to RALF1 (Fig. 3d). The RALF1*–*triggered rapid accumulation of CAR1, 4, 5, 6, 9, and 10 proteins was further confirmed in Myc–tagged CAR proteins (Fig. S7b). Notably, we also tested the accumulation of three CAR proteins (i.e., CAR2, CAR3 and CAR7) which have no interaction with FER, and found that RALF1 induced protein accumulation of CAR2, but not CAR3 or CAR7 (Fig. S7c). This may imply that the RALF peptides and FER modulate CAR protein accumulation in a sophisticated way. In addition, treatment with ABA (10 μM) did not induce the accumulation of CAR9 (Fig. S7d). To answer the RALF1-FER signaling induced CAR protein synthesis by promoting translation rate, we then tested whether the RALF1–FER complex modulated *CAR* mRNA translation using polysome profiling analysis (Zhu *et al*., 2020). From the results, RALF1 promoted the mRNA translation of *CAR1, 4, 5, 6, 9,* and *10* in WT (Fig. 3e). However, the *fer*–*4* mutant blocked RALF1–triggered *CAR1, 4, 5, 6, 9,* and *10* mRNA translation (Fig. 3f), suggesting that RALF1 upregulated the protein synthesis of CAR via FER. Together, these results suggest that RALF1-FER promotes CAR proteins synthesis via facilitating their translation rate, and then the accumulated CAR proteins are recruited to the PM, facilitating the formation of nanoclusters.

### CAR proteins are crucial for RALF1-FER-induced lipid order nanodomains

We next investigated whether RALF1-FER affects nanodomain organizations in the PM. The nanodomain hypothesis propose that lipids in biological membranes may be in a liquid-disordered (Ld) or liquid-ordered (Lo) phase. The Lo phase is enriched in sterols and glycosphingolipids, which are segregated into a more tightly packed phase that is entirely abolished with mβcd treated (Huang *et al*., 2019; Jaillais & Ott, 2020). To test whether RALF1–FER directly impact the lipid order in biological membranes, we took advantage of the lipid order–sensitive probe di–4–ANEPPDHQ (Owen *et al*., 2011; Huang *et al*., 2019; Pan *et al*., 2020). When di–4–ANEPPDHQ molecules detect lipid domains with different dipole potentials in the cell membrane, there is a large shift in the peak emission wavelength of the dye from 630 nm in the Ld phase to 570 nm in the Lo phase (Fig. S8A). The Lo phase shows a higher generalized polarization (GP) value (increased green fluorescence compared with red) (Huang *et al*., 2019). Taking advantage of this assay, we found that *fer*–*4* had a lower GP value than the WT, indicating a general decreased level of ordered lipid nanodomains in the PM (Fig 4a and 4b). Furthermore, treatment with 1 μM RALF1 promoted the higher GP value in WT, but not in *fer-4* (Fig 4a and 4b). To validate this RALF1 effect, we examined lipid order in the presence of MβCD, which depletes sterols, reducing the Lo phase (Huang *et al*., 2019). As expected, the GP value strongly decreased after MβCD treatment in WT (Fig. 4c and 4d), and the effect of RALF1 on trigger Lo phase formation was also suppressed by MβCD treatments (Fig. 4c and 4d).

**Fig. 4.**
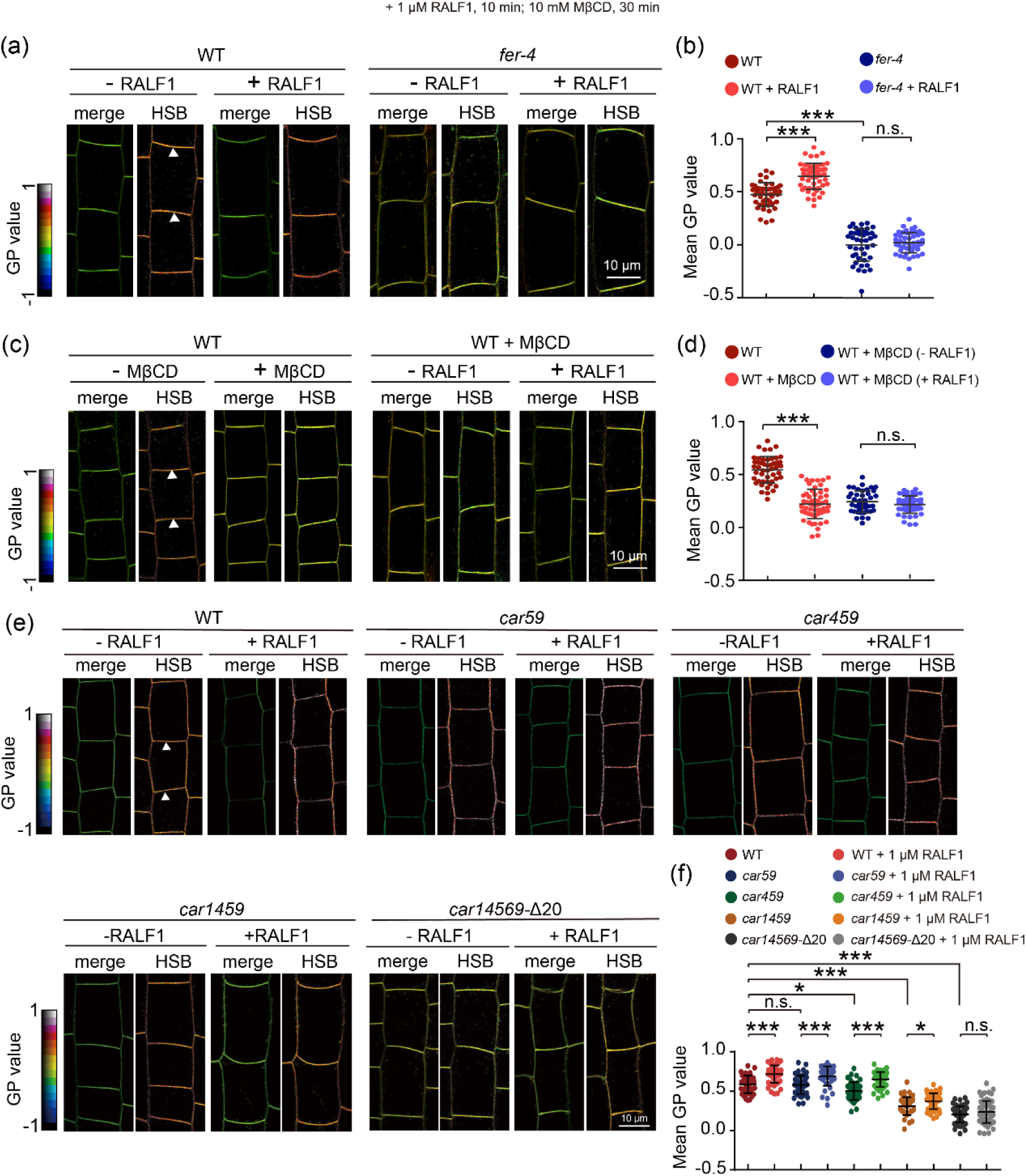
RALF-FER module regulates the formation of ordered lipid nanodomains through CAR proteins. (a) Di–4–ANEPPDHQ lipid staining assay of WT and *fer–4* root cells with or without RALF1 treatment. The GP value is indicated with a colored box. The white triangles indicate the regions for GP value quantification. (b) Statistical analysis of the GP value in (a). The white triangle in (a) indicates the regions for GP value quantification. For each treatment, 42 – 54 cells from 5 roots were measured. (c) Di–4–ANEPPDHQ lipid staining assay of WT root cells with or without MβCD and RALF1 treatments. The GP value is indicated with a colored box. The white triangles indicate the regions for GP value quantification. (d) Statistical analysis of the GP value in (c). The white triangle in (c) indicates the regions for GP value quantification. For each treatment, 42 – 54 cells from 5 roots were measured. (e) Di–4–ANEPPDHQ lipid staining assay in root cells of WT, *car59*, *car459*, *car1459* and *car14569–*Δ20 with or without RALF1 treatment. The GP value is indicated with a colored box. (f) Statistical analysis of the GP value in (e). The white triangle in (e) indicates the regions for GP value quantification. For each treatment, 41 – 60 cells from 5 roots were measured. All experiments were replicated three times with similar results. Data are shown as the mean ± s.d.; *p < 0.05, **p < 0.01, ***p < 0.001, n.s., not significant. One–way ANOVA with Tukey’s test.

We used the Di–4–ANEPPDHQ stain assay to test whether CAR proteins, as the downstream target proteins of RALF1-FER, are directly involved in the formation of ordered lipid nanodomains on the membrane. Thus, the Di–4–ANEPPDHQ stain assay was used to analyze lipid order before and after RALF1 treatment in high–order CAR mutants, which were obtained by different combinations of single and multiple CAR mutants. Amino acid sequences of CARs closely match, and therefore the functions are most likely to be redundant (Rodriguez *et al*., 2014). For example, *car59* double mutant did not show obvious changes in lipid order, and the sensitivity of *car59* to RALF1- triggered higher lipid order formation, was similar to WT using 1 μM RALF1 (Fig. 4e and 4f). We then generated higher–order CAR mutants, and observed a significant difference in lipid order between *car459* or *car1459* mutants and WT (Fig. 4e and 4f). Notably, lipid order of *car1459* showed lower sensitivity to RALF1 than WT (Fig. 4f). Next, we generated 4 independent *car14569* pentuple mutants by designing a CRISPR- Cas9–mediated CAR6 knockout in *car1459* background, termed as *car14569–*Δ1, *car14569–*Δ4, *car14569–*Δ10, *car14569–*Δ20 (all of them harbor out-of-frame deletions in *CAR6*, details in materials and methods). Di–4–ANEPPDHQ stain revealed that the pentuple mutant *car14569–*Δ20 strongly attenuated the lipid order, and as expected, *car14569–*Δ20 displayed insensitivity to RALF1–induced lipid order formation (Fig. 4e and 4f). We also analyzed lipid ordering in WT, *fer*–*4*, and CAR9– Myc/*fer*–*4* plants. When overexpressed in the *fer–4* background, CAR9–Myc partly rescued the lipid order defect of *fer–4* (Fig. S8b and S8c).In addition, to confirm that RALF1 promotion of Lo formation is dependent on the rapid accumulation of CAR proteins, we tested the formation of lipid ordered phase in the *eif4e1 eif(iso)4e* translation initiation factor double mutant (Zhu et al., 2020), which has been identified to be directly involved in the rapid translation of downstream proteins regulated by FER, including CAR proteins. The results showed that membranes exhibited low lipid ordering and no longer responded to RALF1 regulation in the *eif4e1 eif(iso)4e* background (Fig. S9a, S9b, S9e and S9f). Moreover, pretreatment with the translation inhibitor cycloheximide (CHX) before RALF1 indicated that RALF1 no longer promotes Lo phase formation once the rapid accumulation of CAR proteins was prevented by CHX treatment, also showed in *fer-4* (Fig. S9c, S9d, S9e and S9f). Taken together, these data indicate that, rapid accumulation of CAR proteins is necessary for RALF1-FER-triggered higher lipid ordering phase formation, and this process is regulated by multiple CAR family members.

### RALF1-FER-CARs regulates plant immunity by regulating the formation of nanodomains

A recent study found RALF-FER regulates the assembly of immune receptor kinases complexes via modulating plasma membrane nanoscale landscape (Gronnier *et al*., 2022). Our data described above show that the receptor kinase FER regulates membrane nanodomains formation through CAR proteins in response to RALF1 peptide. Thus, we wondered whether the RALF1-FER-CAR axis, through stabilization of the ordered lipid nanodomains at the PM, is able to modulate the assembly of immune complexes. To this end, we first tested the effect of stability of nanodomains on plant responses to RALF and flg22 peptides. ROS burst results showed that when MβCD pretreatment was used to hinder the stability of nanodomains, the response of plants to RALF and flg22 small peptides was significantly weakened, indicating that ordered lipid nanodomains are important for the response to RALF and flg22 peptide signals (Fig. S10a-S10d). To investigate the role of ordered lipid nanodomains stability mediated by FER-CAR in immune regulation, we examined the flg22–induced association between immune receptor kinases FLS2 and its co–receptor BAK1 in *car14569–*Δ20 pentuple mutant. In agreement with previous results (Stegmann *et al*., 2017), we confirmed that the association of FLS2 and BAK1 was severely weakened in *fer-4* compared to WT after flg22 treatment (85% ± 4.5% reduction of the Co–IP signal) (Fig. 5a and 5c). The Co–IP assay of FLS2-BAK1 was also weakened in *car14569–*Δ20 after flg22 treatment compared to WT (25% ± 6.2% reduction), although less dramatically than in *fer–4* possibly because of residual CAR activity remaining in the pentuple *car* mutant (Fig. 5b and 5c). As RALF23 does (Stegmann *et al*., 2017), the activation of FER by RALF1 inhibited FLS2 and BAK1 interaction induced by flg22 in the WT (48% ± 8.2% reduction) (Fig. 5a and 5c). In this co-signaling context, in the absence of FER (*fer-4*), RALF1 had no effect, whereas the residual CAR activity of the *car14569–*Δ20 mutant still mediated the RALF1 inhibitory effect on FLS2-BAK1 interaction, although less efficiently than in WT (compare WT lanes +flg22/+flg22+RALF1 with corresponding *car14569–*Δ20 lanes) (Fig. 5b and 5c). This suggests that a significant portion of RALF1’s inhibitory effect on the flg22 induced association of FLS2–BAK1 is CAR dependent or requires a normal content of CAR proteins.

**Fig. 5.**
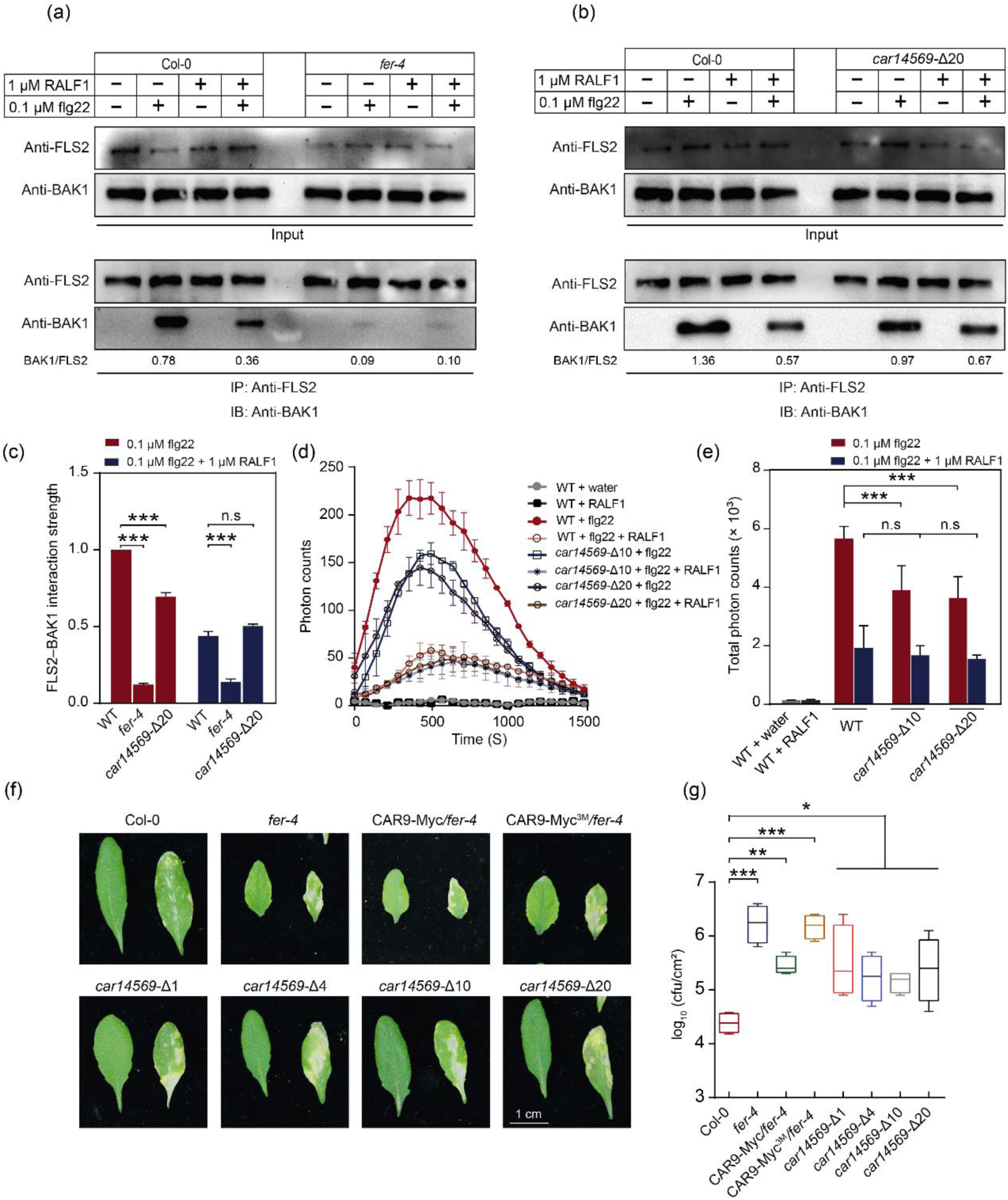
RALF1-FER-CAR axis as a signaling platform to integrate immunity response. (a-b) Co–IP assay showing the weaker interaction between FLS2 and BAK1 in *fer–4* and *car14569–*Δ20 with or without RALF1 and flg22 treatment for 10 min. Western blots were probed with anti–FLS2 and anti–BAK1 antibodies. The relative protein intensity of FLS2 and BAK1 was analyzed using Image J. (c) Quantitative comparison of FLS2 and BAK1 interaction strength in (a) and (b). The ratio of BAK1 to FLS2 in (a) and (b) were normalized. (d-e) The ROS burst was measured after elicitation of leaf discs from the indicated mutant lines with 0.1 μM flg22 and/or 1 μM RALF1 in (d). (e) was showed the ROS integration as mean values of total photon counts over 1500 second in (d). (f) Colony-forming units (cfu) of *Pto* DC3000 bacteria after infection for 3 days. (g) Bacterial populations statistics after infection for 3 days. Bacterial inoculations were performed as described in (f). Log_10_ CFU/cm^2^, log_10_ colony-forming units per cm^2^ of leaf tissue. All experiments were replicated three times with similar results. Data are shown as the mean ± s.d.; ***p < 0.001, n.s., not significant. One–way ANOVA with Tukey’s test.

To further examine the biological relevance of CARs’ function in the RALF–FER pathway, we analyzed CARs’ role in RALF1–FER–mediated immunity response. Flg22–induced ROS production was inhibited in pentuple CAR mutants (Fig. 5d, 5e, S11a, and S11b). The pentuple CAR mutants were more susceptible to the *Pseudomonas syringae pv.tomato* (*Pto*) DC3000, albeit not to the same extent as *fer-4* (Fig. 5f and 5g). In addition, overexpression of wild-type CAR9 in *fer-4* partially restored the immunodeficient phenotype of *fer-4*, but not CAR9^3M^ (Fig. 5f and 5g). Taken together, the formation of ordered lipid membrane nanodomains regulated by RALF1-FER-CAR module is necessary for the regulation of plant immunity, which depends on the assembly of immune receptor complexes in the regulatory membrane nanodomains.

In addition, FER has also been reported to be involved in cell growth, hormone signaling, stress responses (Zhu *et al*., 2021). Notably, by comparing the phenotype of *fer-4* and multiple CAR mutants, we observed that *fer-4* and CAR mutants showed a similar defect in root hair and epidermal cell growth and overexpression of CAR9 in *fer-4* background was also able to partly rescue the root hair growth defect (Fig. S12a- S12d). Further analysis the effects of temperature stresses on *fer-4* and *car14569–*Δ20, similarly, both mutants showed hypersensitive phenotypes to temperature stress (Fig. S12e-S12h). Together, this set of results shows that RALF1 induces the formation of ordered lipid nanodomains through CAR proteins, which is crucial for FER to regulate multiple signaling pathways.

## Discussion

The molecular mechanisms through which eukaryotic cells modulate their PM landscape via activating receptor kinases in response to external signals remain unclear (Jaillais & Ott, 2020). Our study reveals that FER regulates the general lipid order nanodomains of the membrane in response to extracellular peptides, RALFs. Proteins with C2 domain have been shown to be involved in protein transport and membrane fusion via interacting with negatively charged phospholipid (Martens *et al*., 2007; Martens & McMahon, 2008; Rodriguez *et al*., 2014). A lipid binding protein family with C2 domain, CAR, facilitates the ordered lipid nanodomains formation via rapid accumulation of protein in response to RALF1 (within 10 min after RALF1 treatment). Multiple CAR proteins interact with FER at the PM and are phosphorylated by the kinase (20 min after RALF1 treatment), thus weakening the ability of CARs to interact with lipids to form a negatively feedback. The higher ordered lipids, which are particularly induced by RALF1-FER-CAR axis, might be involved in signaling multiple rapid responses, including immunity (Fig. 6).

**Fig. 6.**
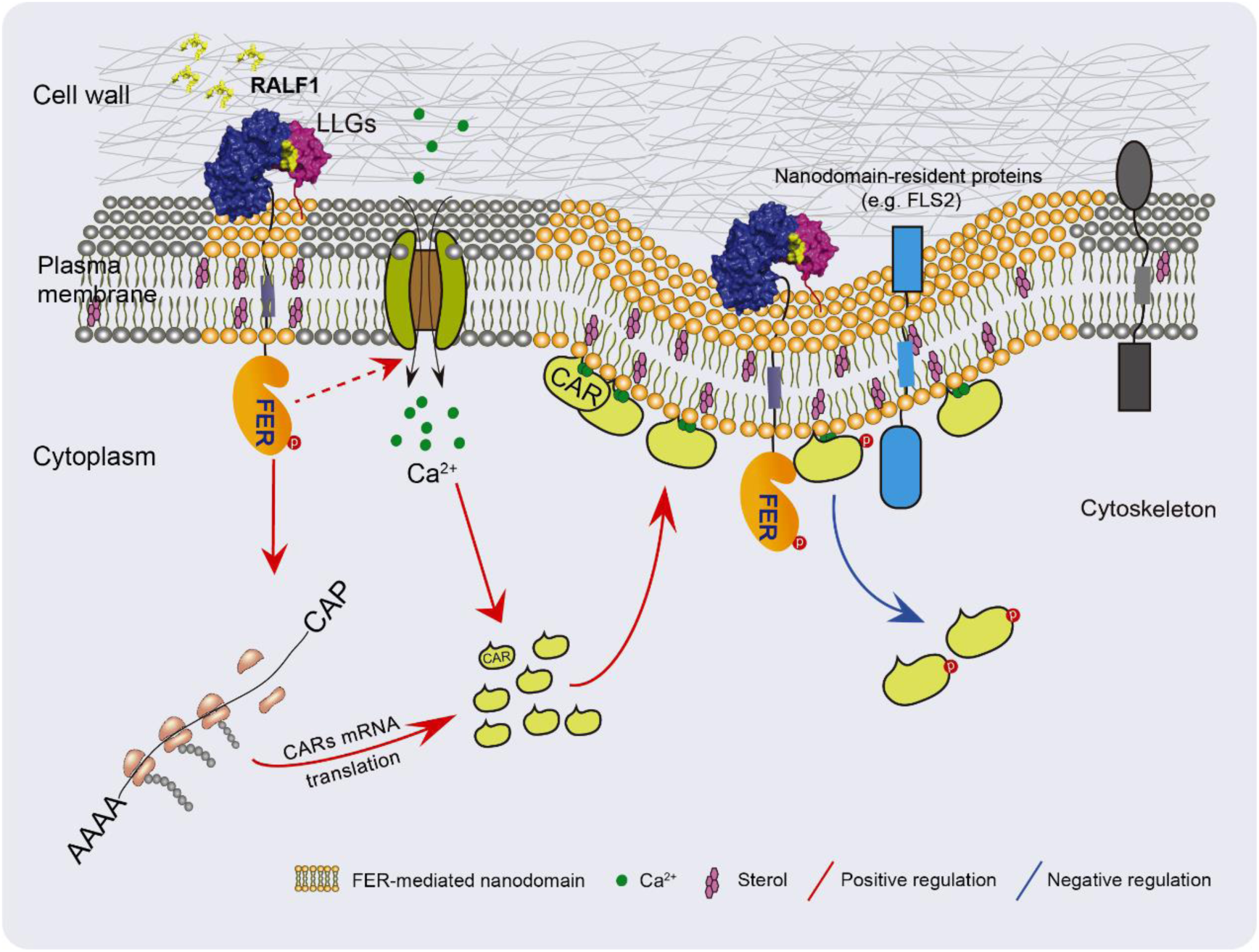
A working model for RALF1-dependent plant response through the FER- CAR axis in the formation of proper lipid ordered nanodomains. The RALF1–FER signaling first promotes CAR protein synthesis, and then, more CAR proteins localize to the PM via binding Ca^2+^ and interacting with negative charge phospholipids, which positively regulates formation of lipid ordered nanodomains and membrane curvature. Finally, FER interacts with and phosphorylates CARs in a RALF1–dependent manner. The enhanced phosphorylation level of CARs reduces their binding to negatively charged phospholipids. This maintains the dynamic balance of CAR proteins on the membrane, thus, serving for fine-tuning of the system sensitivity. Proteins and structures are not to scale.

An early and rapid response to the RALF1 peptide is the FER–triggered Ca^2+^ wave (Ngo *et al*., 2014). Thus, it is reasonable to assume that the RALF1–FER–induced Ca^2+^ wave can further enhance the PM location of CAR proteins (Rodriguez *et al*., 2014). In agreement with previous studies, we found that Ca^2+^ upregulates the lipid binding and membrane curvature ability of CARs (Rodriguez *et al*., 2014; Diaz *et al*., 2016), which opens the possibility that RALF1 promotes the rapid accumulation of CAR proteins by activating FER and further induces membrane curvature through interaction of CARs with anionic lipids. Furthermore, we found that FER-dependent phosphorylation of CARs, which occurs in the proximity of conserved Asp residues involved in Ca^2+^ binding (Diaz *et al*., 2016), seems to impair CAR binding to vesicles. Therefore, in the first minutes following RALF1 perception (<10 min), FER stimulates CAR proteins accumulation by upregulating its translation and might also trigger CAR membrane recruitment in a Ca^2+^-dependent manner. In a second step (>20 min), the C2 domain of CARs by phosphorylation of specific Ser/Thr/Tyr residues might tune CAR binding to membranes. Mechanistically, the addition of negative charges within the lipid-binding pocket of the C2 domain by phosphorylation might repel the interaction with negatively charged phospholipid in the PM (Simon *et al*., 2016; Platre *et al*., 2018). Thus, the system appears to be designed for a fast-self-amplified recruitment of CAR in PM, followed by a brake mechanism leading to CAR membrane dissociation. This scenario becomes even more complex when considering that RALF1–FER signaling is a master regulator of cell wall integrity (Zhang *et al*., 2020), and the cell wall also has profound roles in the diffusion coefficients and PM compartmentalization of resident proteins (Luu *et al*., 2012; Martiniere *et al*., 2012; McKenna *et al*., 2019; Danek *et al*., 2020; Jaillais & Ott, 2020).

A recent study unraveled the regulation of FLS2 and BAK1 nanoscale organization by the RALF receptor FER (Gronnier *et al*., 2022). How FLS2 and BAK1 associate in a ligand-dependent manner within the plasma membrane in response to RALF peptides needs further study. Herein, we found that the receptor kinase FER mediated formation of proper lipid order nanodomains via CAR proteins, which may offer a strategy to modulate the formation of other receptor kinase complexes (e.g., FLS2 and BAK1)to interact with their downstream effectors; therefore, FER could promote the integration of multiple environmental cues. Various studies indicate that FER regulates many aspects of plant biology, including growth, development, environmental adaptability, and immunity, by sensing a variety of external signals (Zhang *et al*., 2020; Zhu *et al*., 2021). Similarly, we also found a pleiotropic phenotype in the high–order CAR mutants (Rodriguez *et al*., 2014), indicating that CAR-mediated regulation of membrane order could be pivotal for additional signaling systems.

Interesting future research topics include whether RALF1–FER regulates various cell type–specific outputs and/or signaling events via the dynamic nanoclustering of proteins and lipids (involving other receptor kinase complexes, ROPs, CARs and phosphatidylserine) of the PM. More studies are also needed to verify whether signal– triggered PM compartmentalization based on kinase receptors and C2 domain– containing proteins is a common mechanism in eukaryotic cells. In addition, previous evidence indicates that CAR proteins are located not only in the PM but also in the cytoplasm and nucleus, where these proteins participate in ABA signaling (Rodriguez *et al*., 2014) and interact with LOT1 to enhance plant drought tolerance (Qin *et al*., 2019). It would be interested to address whether the RALF1-mediated regulation of CAR accumulation, its recruitment membrane interaction dynamics, and its phosphorylation may also impact ABA signaling.

## Data availability

The authors declare that the data supporting the findings of this study are available within the paper and its supplementary information.

## Acknowledgments

We thank Liming Xiong, Tao Qin, and Chao Li for providing plant materials and Cyril Zipfel for critical comments and suggestions. This work was supported by grants from National Natural Science Foundation of China (NSFC–31871396, 31571444), Young Elite Scientist Sponsorship program of CAST (YESS20160001), and the Open Research Fund of the State Key Laboratory of Hybrid Rice (Hunan Hybrid Rice Research Center) to F.Y. China Postdoctoral Science Foundation funded project (2020M672475), and the Science and Technology Innovation Program of Hunan Province (2021JJ40060) to F.X. Work in P.L.R. laboratory was supported by grant PID2020-113100RB funded by MCIN/AEI/ 10.13039/501100011033.

## Author contributions

F.Y., and W.C. conceived the project and designed the research; W.C., H.Z., S.Z., F.X., M.Y., G.X., J.C. A.C., L.R., Y.L, Q.X., and Q.F performed the research; D.W., X.L., A.C., X.L, and P.L.R. contributed new reagents/analytic tools; F.Y. and W.C. analyzed data and wrote the paper with contribution from P.L.R. and Y. J.; all authors reviewed and approved the manuscript for publication.

## Supporting information

**Fig. S1.**
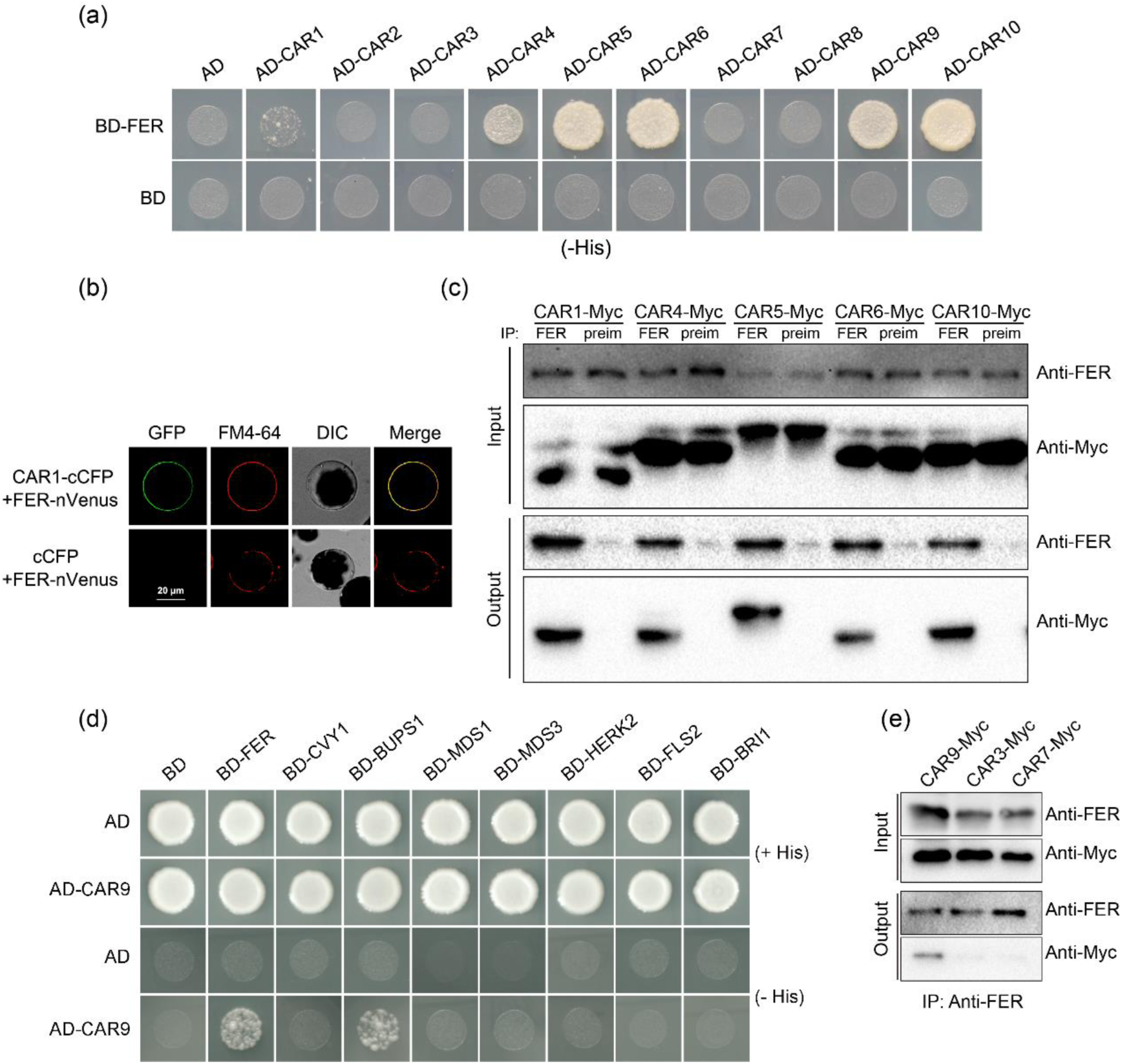
FER physically interacts with CARs. (a) Y2H assays showing the interaction between CARs and FER. SD III (–His/–Leu/– Trp) selection medium containing 20 mM 3–AT was used to test the interaction. The CAR members were cloned into the AD vector and the FER–CD was cloned into the BD vector. (b) FER interacts with CAR1 in BiFC assays in *Arabidopsis* protoplasts. The negative control (cCFP + FER–Venus) is shown, bar = 20 μm. (c) Co–IP assays. The immunoprecipitated FER and coimmunoprecipitated CARs were indicated using anti–FER and anti–Myc antibodies, respectively. (d) Y2H analysis of interaction between CAR9 and multiple CrRLK1L subfamily members, FLS2 and BRI1. SD/– Ade/–Leu/–His selection medium containing 20 mM 3–AT was used for screening yeast growth. The CAR9 was cloned into the AD vector. The cytoplasmic domain of CrRLK1L subfamily members, FLS2 and BRI1 were cloned into the BD vector. (e) Co-IP assays. The immunoprecipitated FER and coimmunoprecipitated CARs were indicated using anti–FER and anti–Myc antibodies, respectively. At least three biological replicates of (a) – (e) were performed with similar results.

**Fig. S2.**
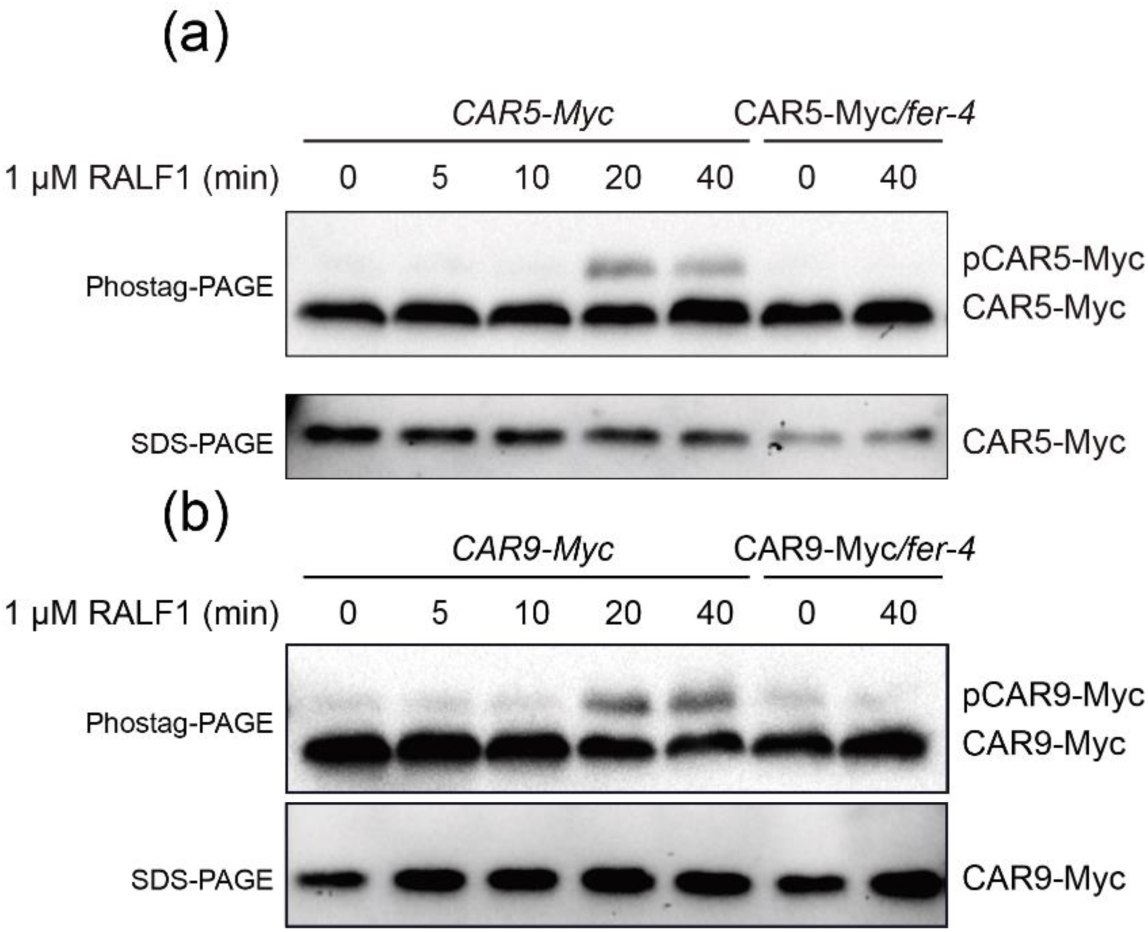
Phostag–PAGE analysis dynamics of CAR5 and CAR9 protein phosphorylation. (a-b) Phosphorylated CAR proteins were assay in *CAR*–*Myc* and CAR–Myc/*fer*–*4* seedlings. Phosphorylated CAR–Myc bands are indicated as pCAR– Myc. Three biological replicates were performed with similar results.

**Fig. S3.**
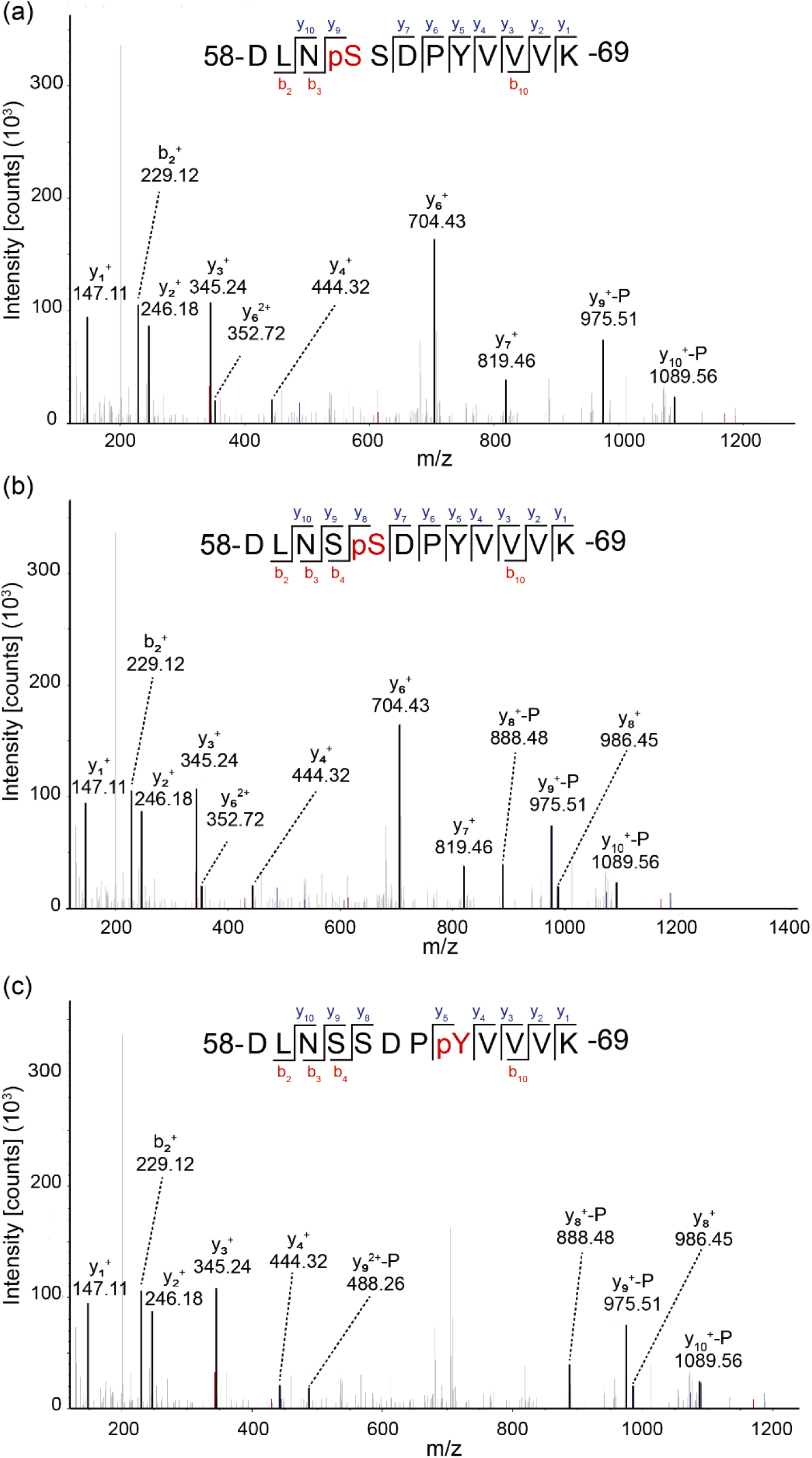
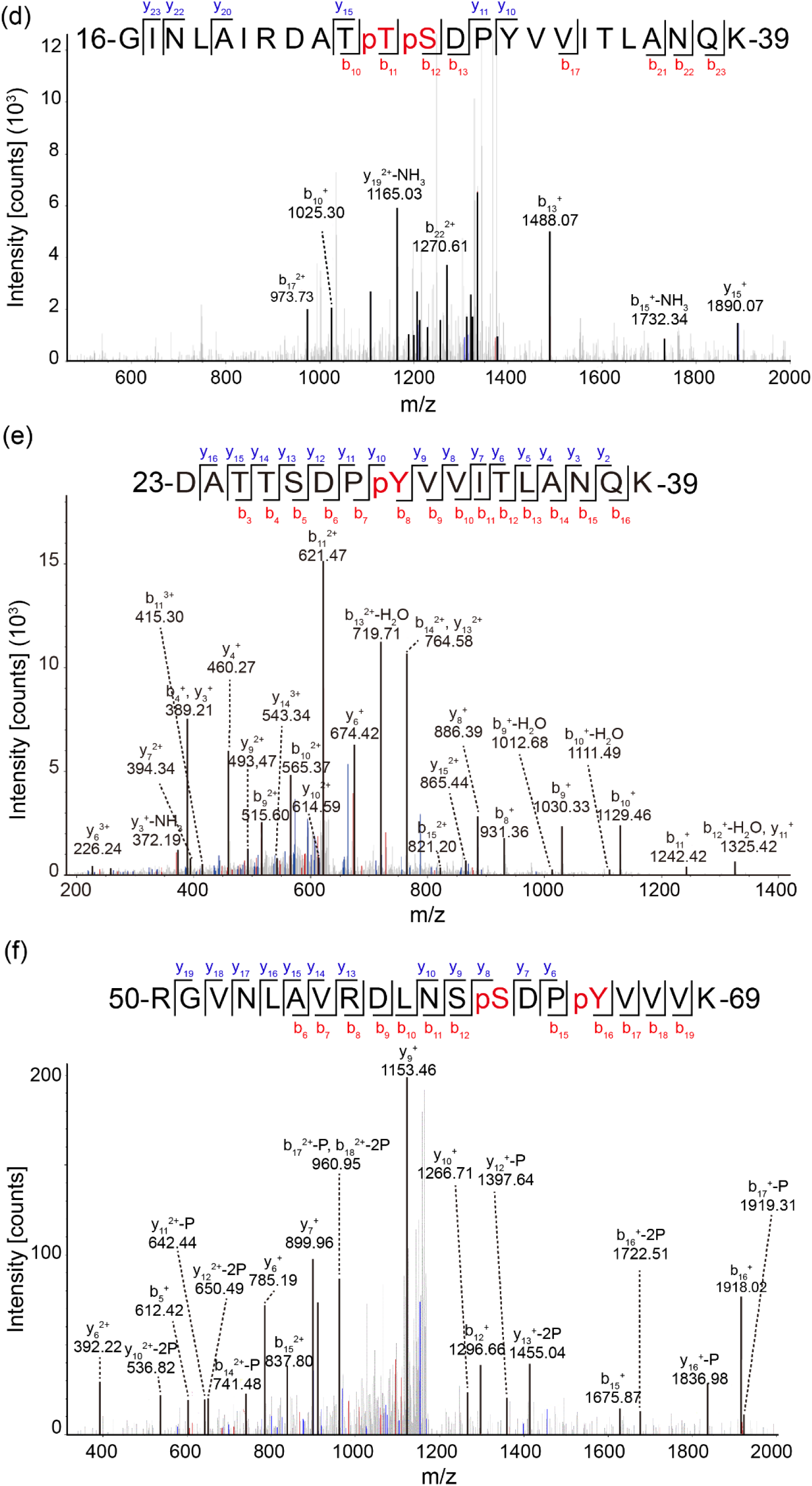
ESI mass spectrometry identification of CARs phosphorylation sites. (a-c) The identified CAR5 phosphorylation sites of Ser61, Ser62, and Tyr65 in *E. coli* coexpression assays. (d-e) The identified CAR9 phosphorylation sites of Thr26, Ser27, and Tyr30 in *E. coli* coexpression assays. (f) The identified IP CAR5–Myc phosphorylation sites of Ser62 and Tyr65 from plant *in vivo* proteins. The identified peptide sequences are shown. The y–ion and b–ion is shown in the sequences.

**Fig. S4.**
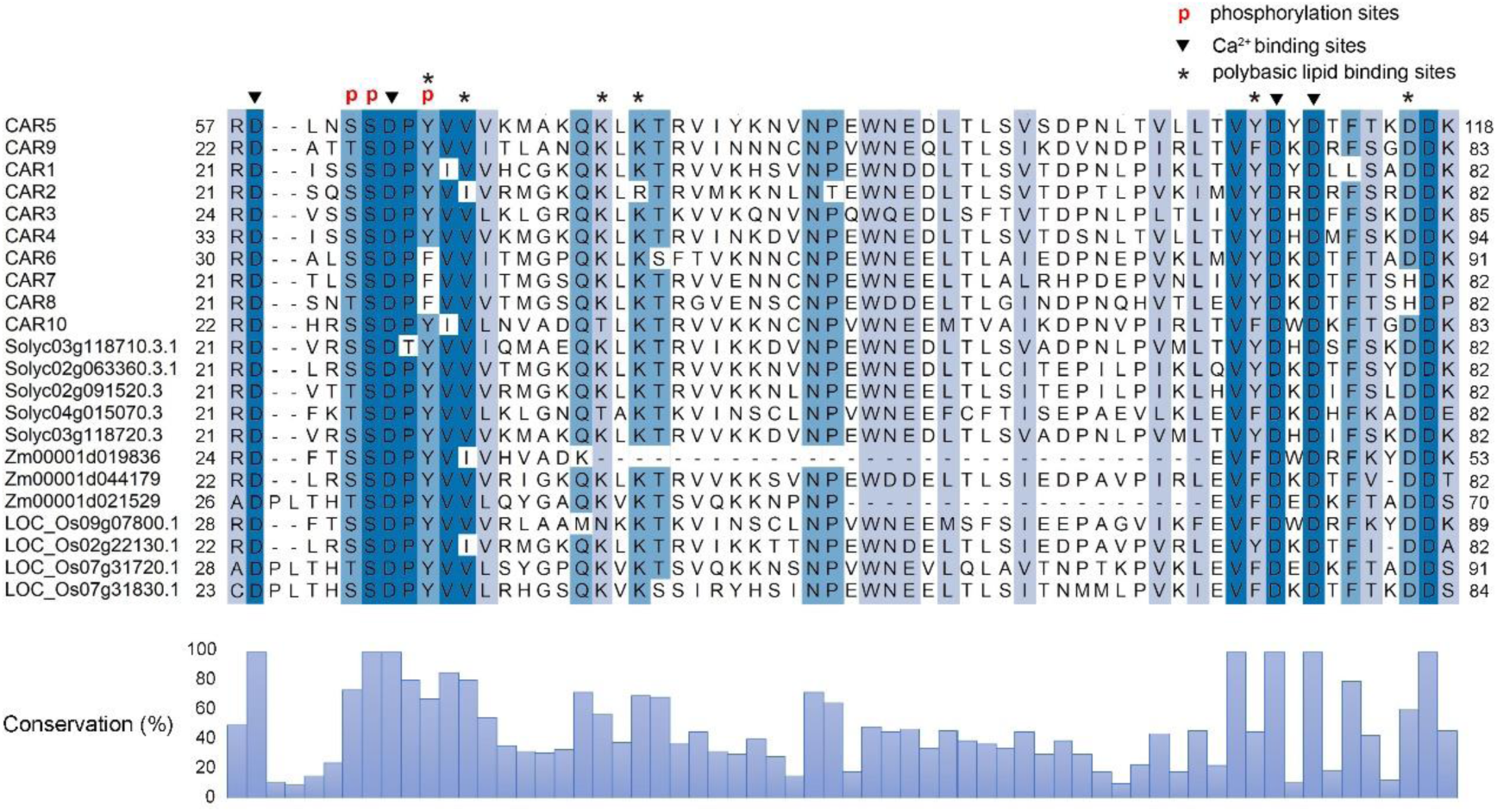
FER triggered phosphorylation sites of CAR members. The alignment of CAR homologs from *Arabidopsis thaliana*, tomato (*Solanum lycopersicum*), maize (*Zea mays*), and rice (*Oryza sativa*). Identical residues are highlighted with a blue background. The identified *Arabidopsis* CAR protein phosphorylation sites regulated by FER are indicated in the sequence with the letter p. The residues essential to Ca^2+^ binding is marked with triangles. The residues essential to lipid binding are marked with asterisks. The data were generated using Clustal X and BioEdit.

**Fig. S5.**
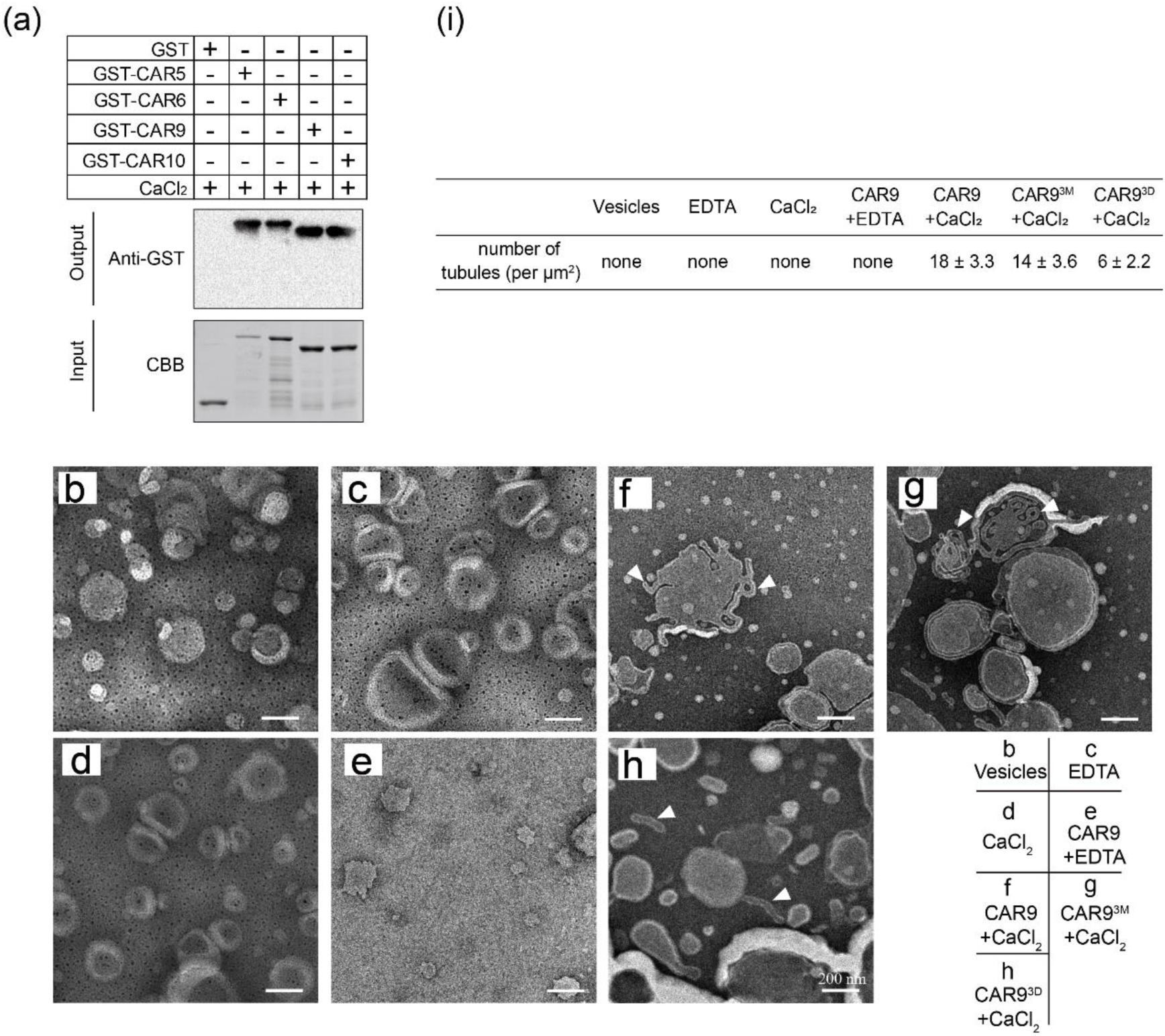
CAR proteins directly interact with anionic lipids and promote lipid tubules. (a) Lipid binding abilities of CAR5, 6, 9, and 10 proteins (0.1 μM) in the presence of 1 mM CaCl_2_. (b-h) Negative stain transmission electron micrographs. Negative-stain transmission electron micrographs of 12.5 μM liposomes incubated with 0.1 μM CAR9 plus 1 mM CaCl_2_. The white triangle indicates the lipid tubules induced by CAR proteins. The average diameter of tubules was approximately 17.4 ± 3.4 nm from the outer bilayer to the outer bilayer. (i) Lipid tubules quantity analysis in (b-h). The quantity was counted as the number of lipid tubules per μm^2^. At least three biological replicates were performed with similar results. bar = 200 nm. All experiments were replicated three times with similar results.

**Fig. S6.**
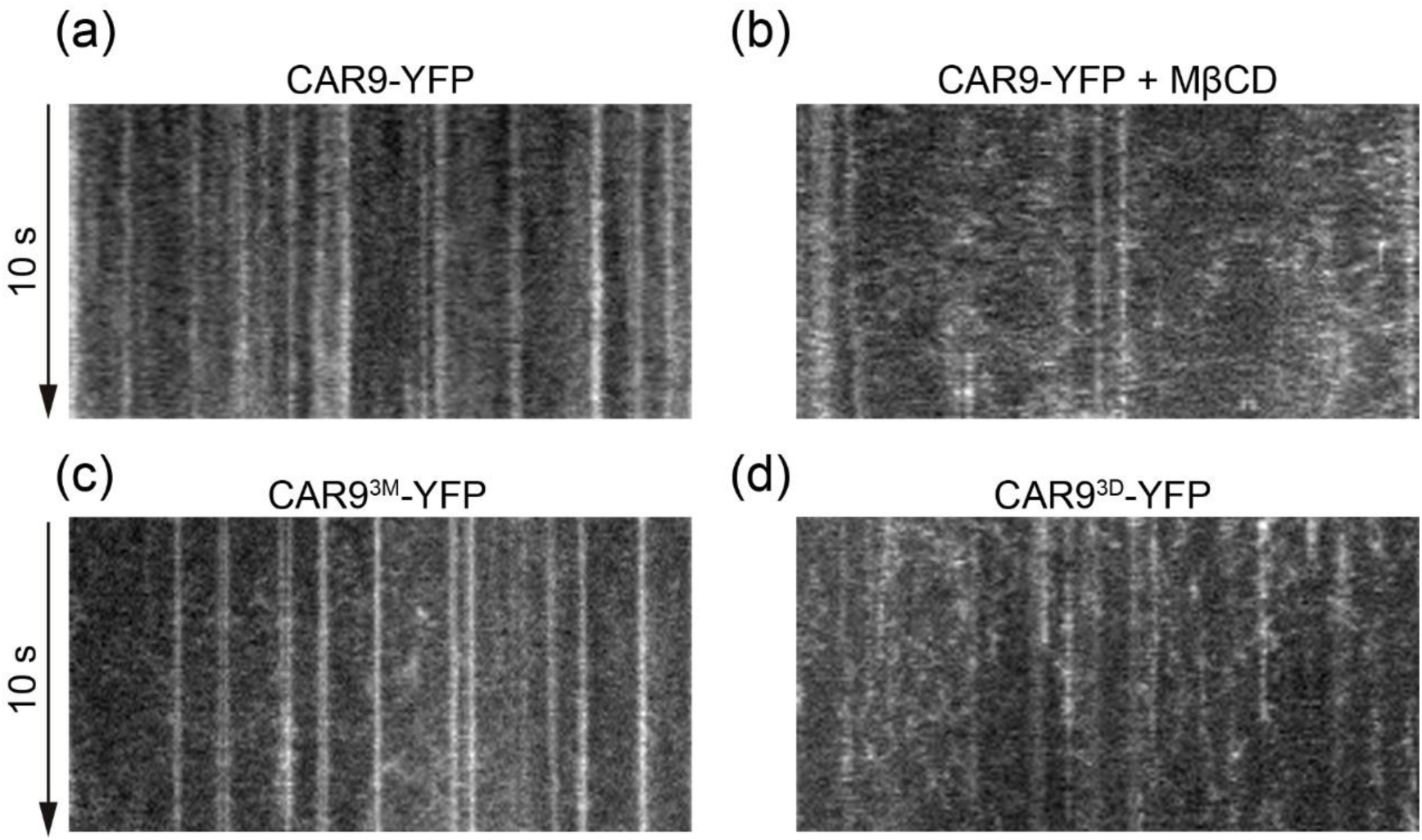
Kymograph analyze the effect of phosphorylation on the dwell time of CAR9 in nanodomain. (a-d) Representative kymograph showing the dwell times of CAR9 (a), CAR9 + MβCD (b), CAR9^3M^ (c), CAR9^3D^ (d). Data were analyzed by Image J.

**Fig. S7.**
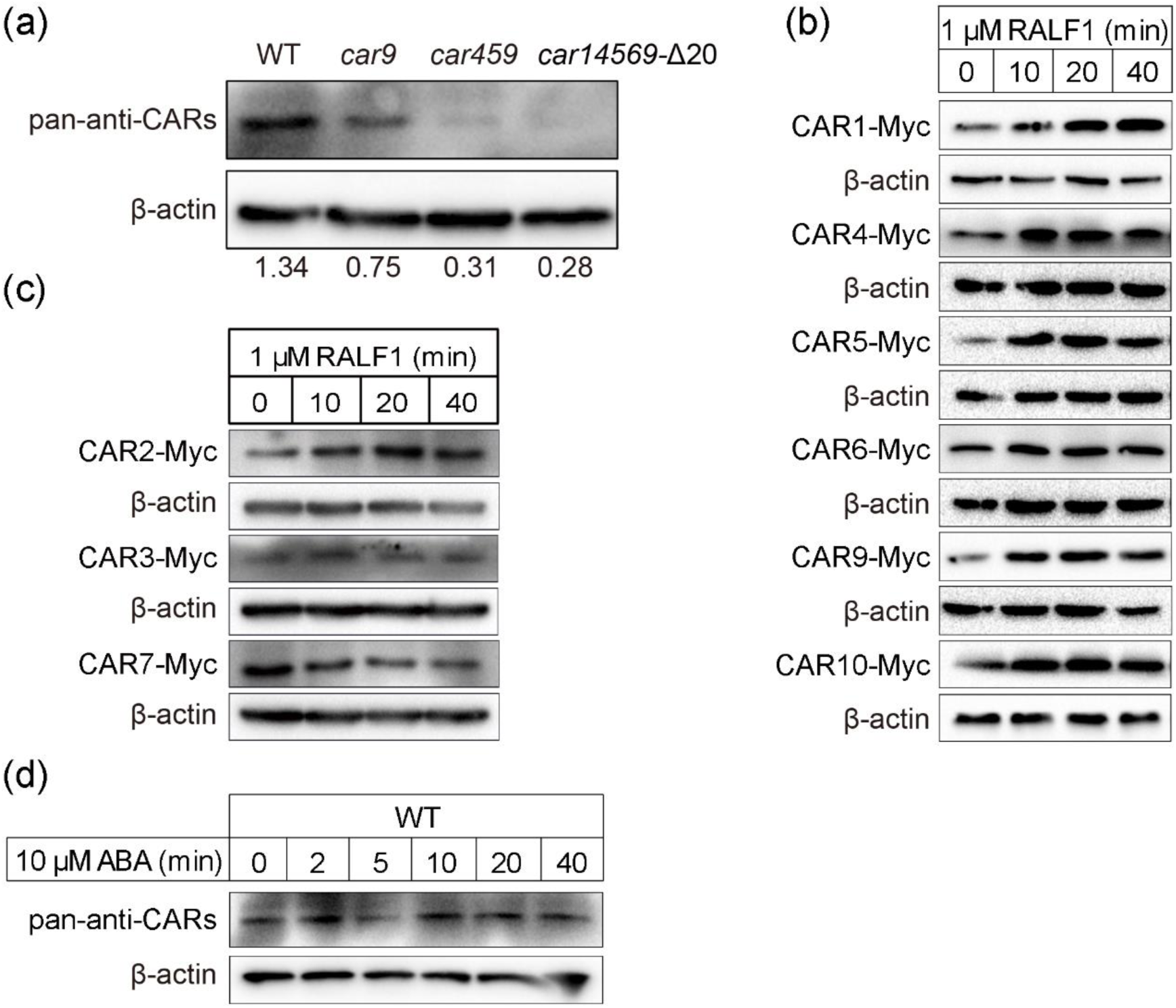
RALF1 induces CARs protein accumulation. (a) Western blot detection of pan–anti–CARs antibody in WT and respective mutants. (b-c) CAR proteins accumulation was analyzed following treatment with RALF1 in the corresponding transgenic lines. β–actin acted as the loading control. (d) CAR proteins accumulation was analyzed following treatment with ABA in WT using pan–anti–CARs antibody.

**Fig. S8.**
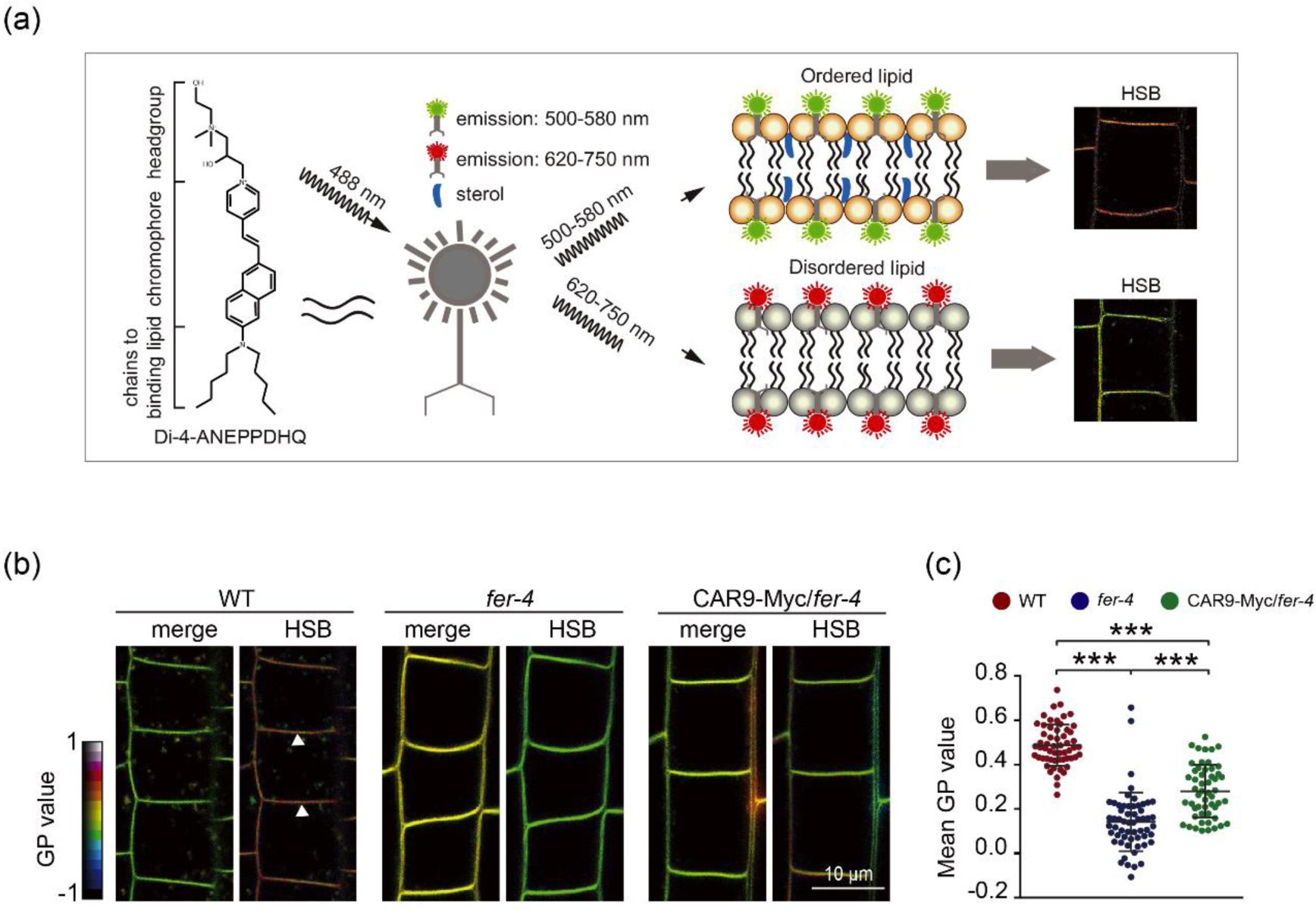
CAR9 partially rescued the lipid order of *fer-4*. (a) Schematic diagram of di–4–ANEPPDHQ staining (HSB: hue-saturation-brightness). (b) Di–4–ANEPPDHQ lipid staining assay of WT, *fer–4*, and CAR9–Myc/*fer–4*. The GP value was indicated with a colored box. (c) Statistical analysis of the GP value in (b). The white triangle in (b) indicated the regions for GP value quantification. For each treatment, 41–50 cells from 5 roots were measured. At least three biological replicates were performed with similar results. Data are shown as the mean ± s.d., ***p < 0.001. One–way ANOVA with Tukey’s test.

**Fig. S9.**
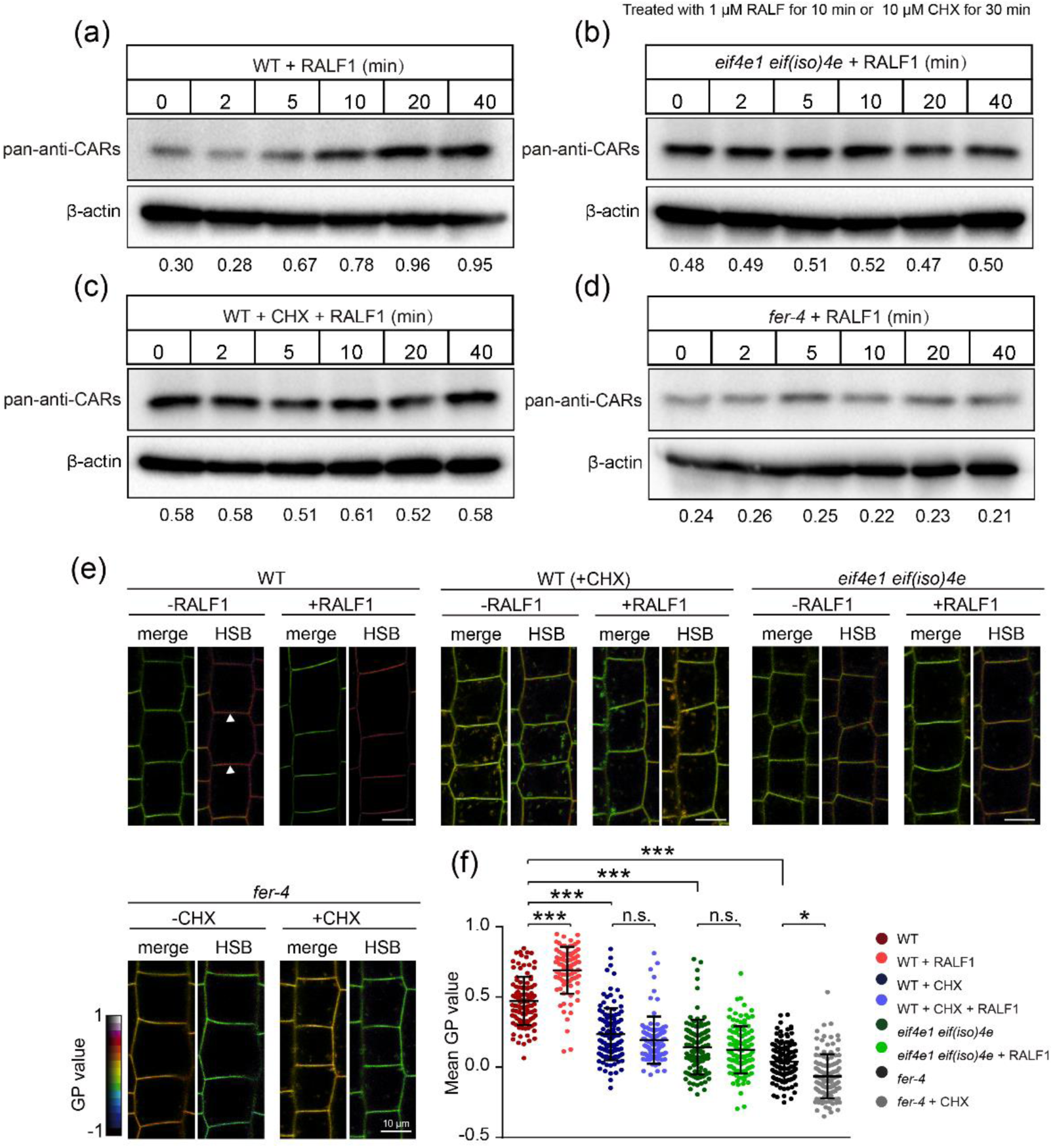
RALF1 induces the formation of lipid ordering through rapid accumulation of CAR proteins. (a) RALF1-induced CAR proteins accumulation in WT. CAR proteins were detected by pan-anti-CARs antibody. The relative CAR proteins accumulation was analyzed using ImageJ. (b) RALF1-induced CAR proteins accumulation was blocked in *eif4e1 eif(iso)4e*. (c) CHX blocked RALF1-induced rapid accumulation of CAR proteins. Seedlings were pretreated with 10 μM CHX for 30 min before RALF1 treatment. (d) RALF1-induced CAR proteins accumulation was blocked in *fer-4* mutant. (e) Di–4–ANEPPDHQ lipid staining assay of WT, *eif4e1 eif(iso)4e, fer–4* root cells with or without RALF1 and CHX treatment. (f) Statistical analysis of the GP value in (e). The white triangle in (e) indicates the regions for GP value quantification. At least three biological replicates were performed with similar results. Data are shown as the mean ± s.d., * p < 0.05, ***p < 0.001. n.s., no significant. One–way ANOVA with Tukey’s test.

**Fig. S10.**
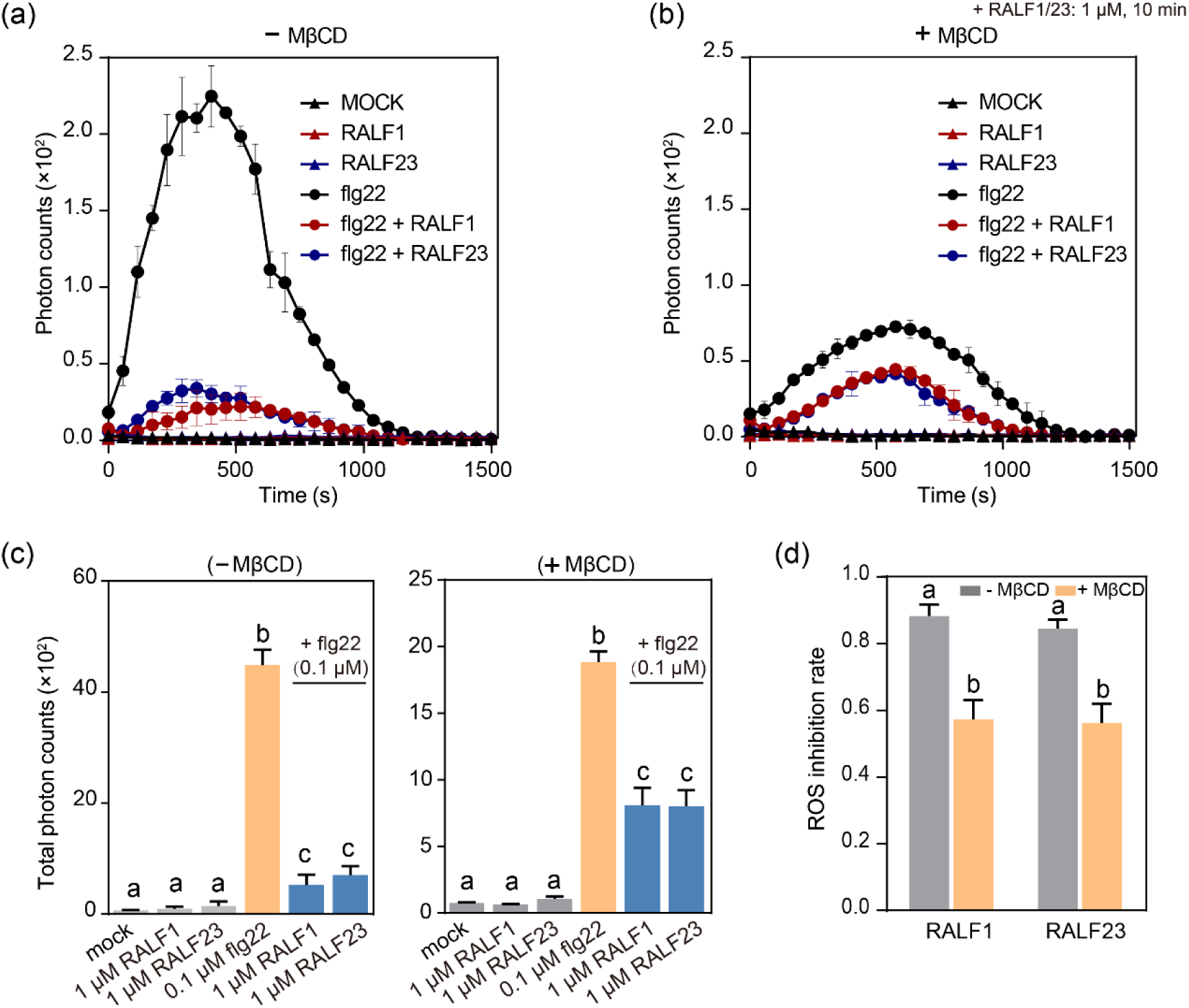
Lipid order nanodomains are crucial for plant immunity. (a) ROS burst progression in Col–0 leaf discs treated with flg22, RALF1, or RALF23. For each treatment, four leaves from 4–week–old plants were measured within 25 min. (b) ROS burst progression in MβCD–pretreated Col–0 leaf discs treated with flg22, RALF1, or RALF23, as indicated. For each treatment, four leaves from 4–week–old plants were measured within 25 min. (c) The values of total photon count over 25 min in (a) and (b). (d) Inhibition rate of ROS production in (a) and (b). MβCD reduced the inhibition of RALF1 and RALF23 treatment to flg22–induced ROS production. All experiments were replicated three times with similar results. Data are shown as the mean ± s.d.; different letters stand for P < 0.05. One–way ANOVA with Tukey’s test.

**Fig. S11.**
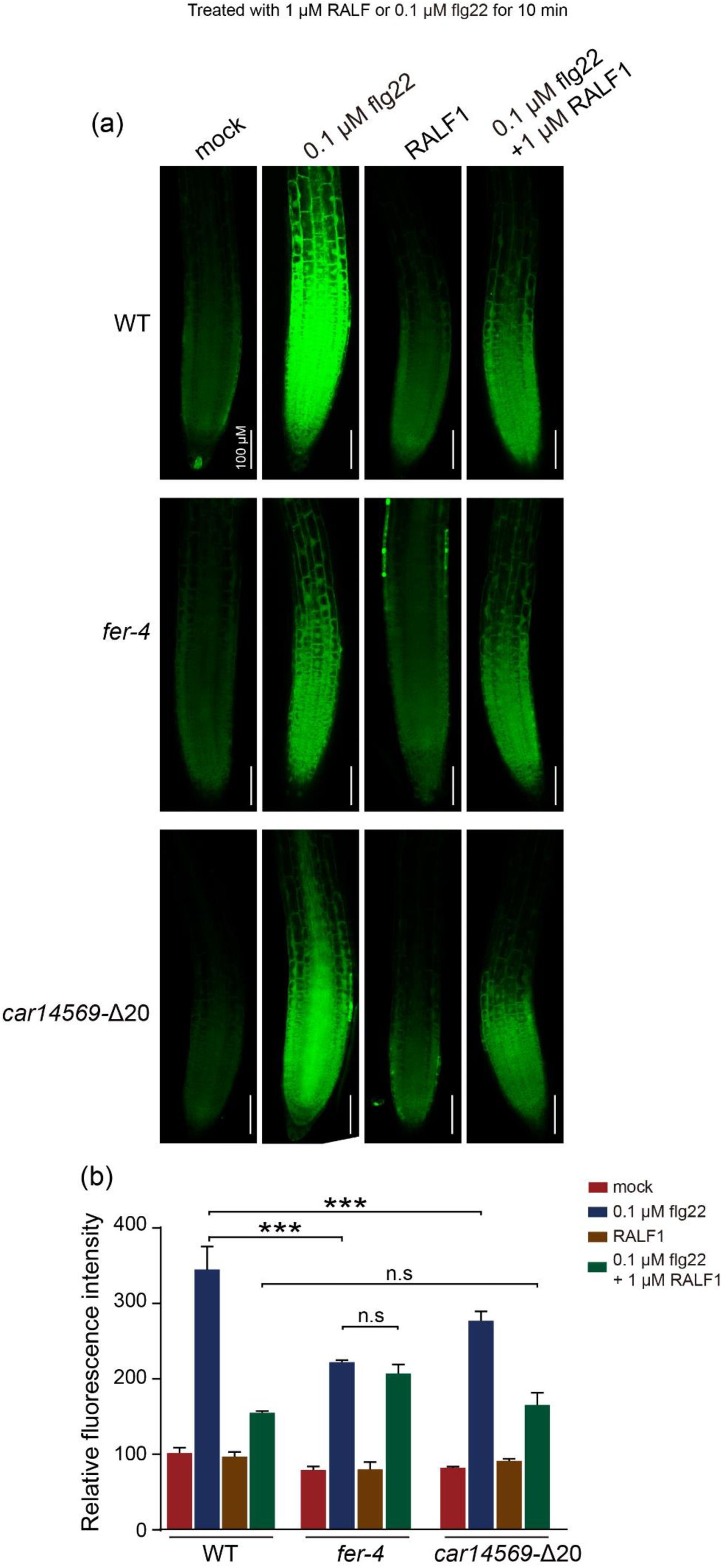
FER-CAR module is crucial for immune response. (a) Representative images of H2DCF-DA-stained roots of WT (Col-0), *fer-4*, and *car14569–*D20 treated with flg22 and/or RALF1 for 10 min. (b) Quantified average H2DCF-DA signal intensity in (a). 5-10 roots from each group were measured. At least three biological replicates were performed with similar results. Data are shown as the mean ± s.d.; ***, p < 0.001, and n.s., not significant. One–way ANOVA with Tukey’s test.

**Fig. S12.**
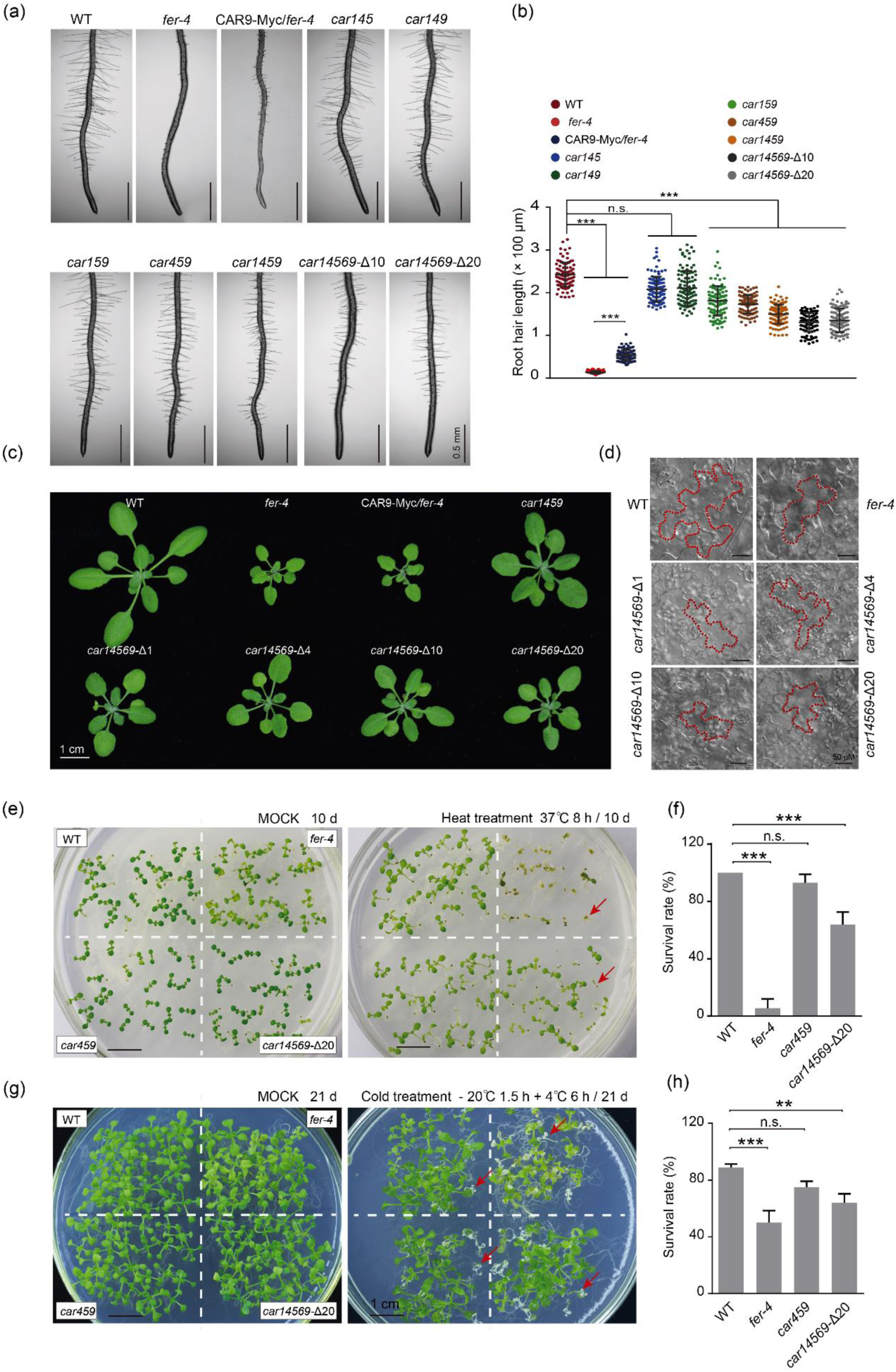
*fer-4* and multiple CAR mutants phenotype analysis. (a) The root hair phenotype of multiple CAR mutants. Seedlings grown vertically on solid 1/2 MS medium for 4 days for root hair phenotype observation. (b) Root hair length statistics in (a). More than 100 root hair of each phenotype were measured. (c) Vegetative growth phenotypes analysis of CAR mutants, *fer–4* and CAR9–Myc/*fer–4*. Observing the vegetable growth phenotypes after the seedlings grow in the soil for about 3 weeks. (d) Pavement cell (PC) phenotypes of 6-day-old seedlings. (e) Heat stress treatment assays. *car459* and *car14569–Δ*20 showed higher sensitivity to heat treatments than the WT. The red arrows indicate the dead seedlings. (f) Statistical analysis of survival rate in (e). 72 seedlings of each phenotype were measured. (g) Cold stress treatment assays. *car459* and *car14569*–*Δ*20 showed higher sensitivity to cold treatments than the WT. The red arrows indicate the dead seedlings. (h) Statistical analysis of survival rate in (g). 72 seedlings of each phenotype were measured. All experiments were replicated three times with similar results. Data are shown as the mean ± s.d.; *p < 0.05, **p < 0.01, ***p < 0.001, n.s., not significant. One–way ANOVA with Tukey’s test.

**Table S1.**
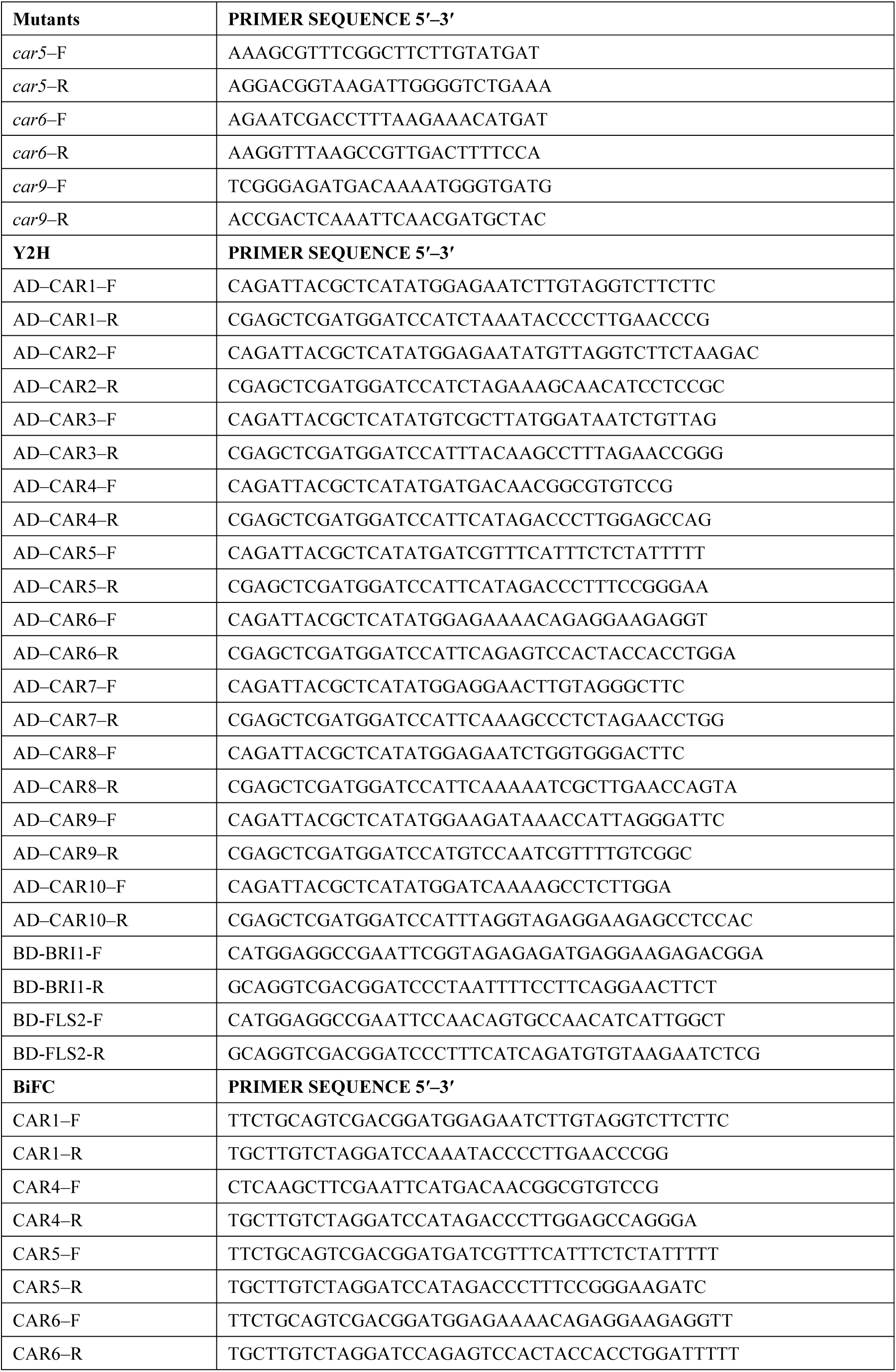
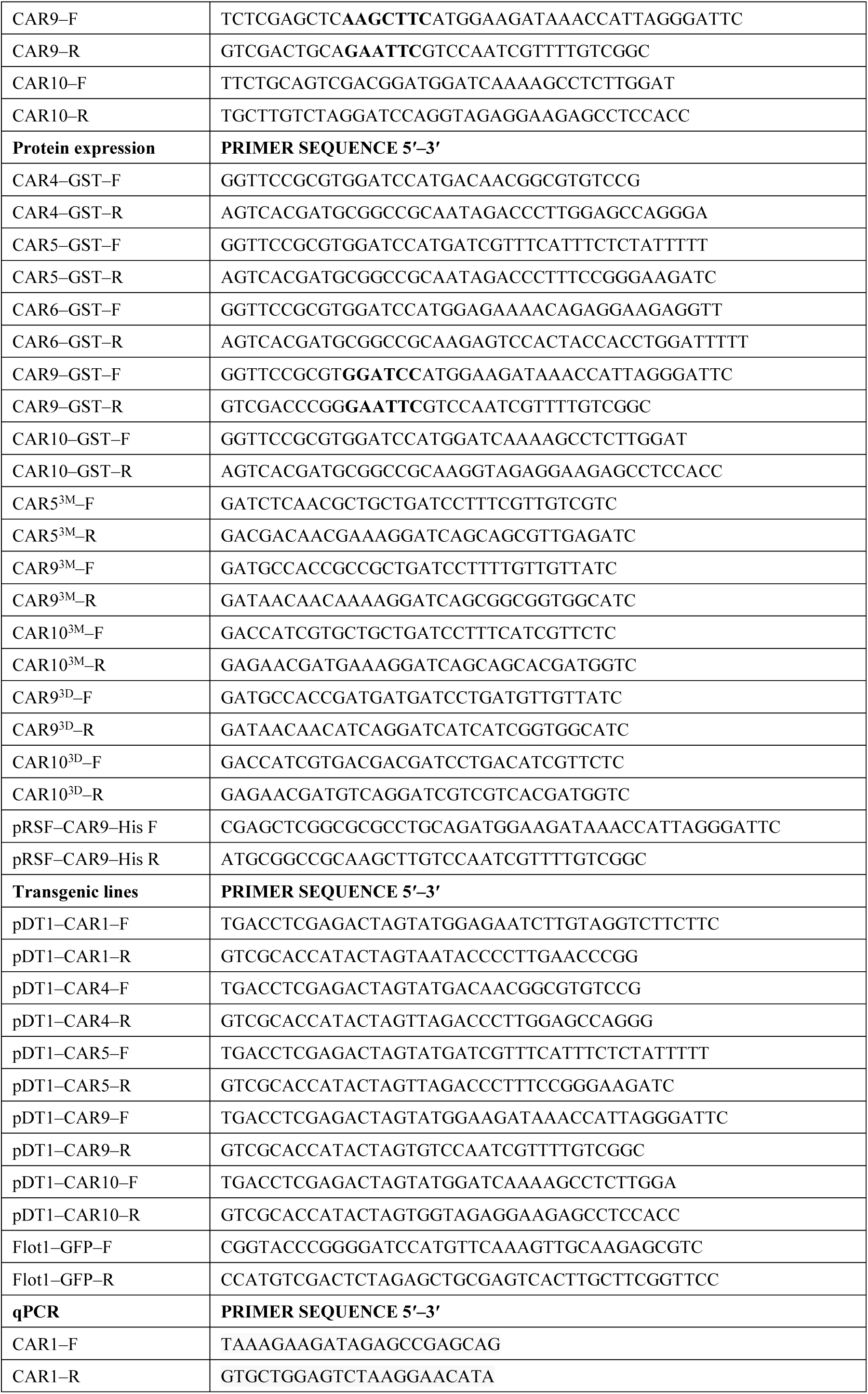
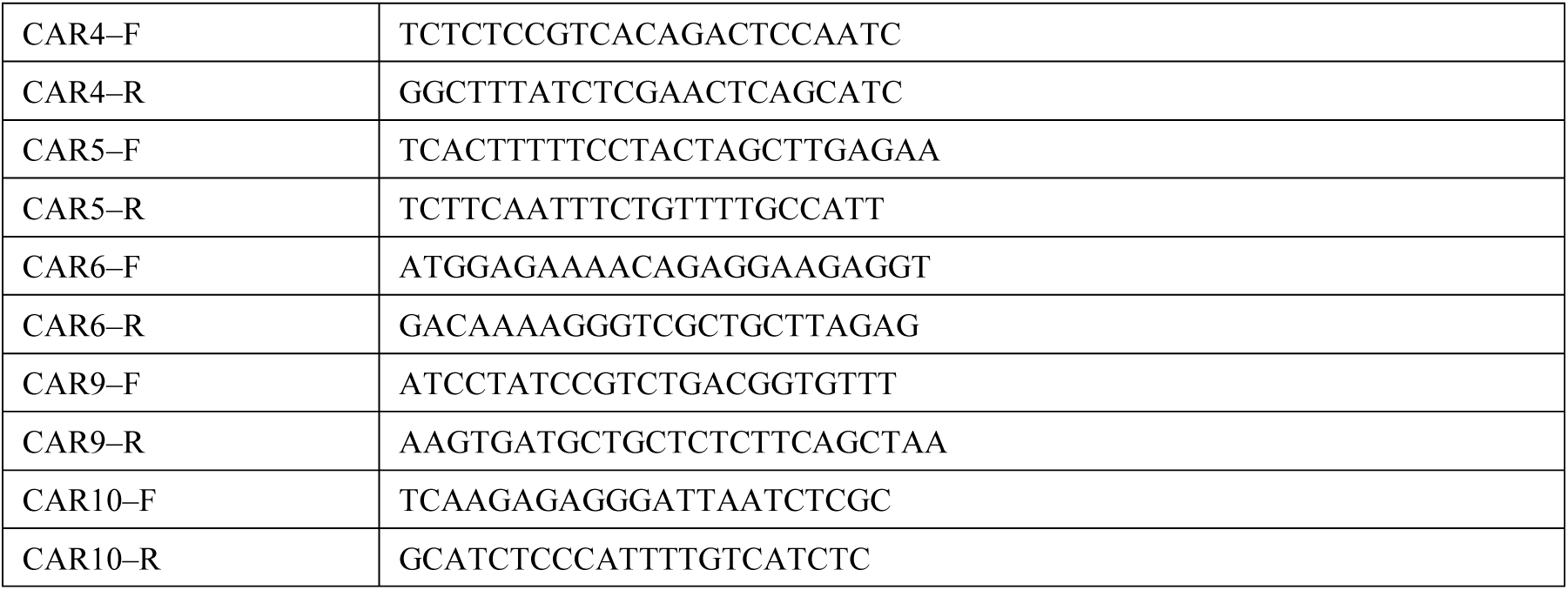
Primers used in this study.

## Supporting information methods S1

### Plant Material and Growth Conditions

*Arabidopsis thaliana* seeds were first surface sterilized by 75 % alcohol. After stigmatization at 4 °C for 3 d, seeds were growing on 1/2 MS with 0.8 % sucrose and 1.0 % Phytagel (Sigma–Aldrich) at 22 °C in a 16–h light/8–h dark condition for subsequent analysis. The *car5* (SAIL_802_B08), *car6* (SALK_052572.54.00.x), *car9* (SALK_088115.56.00.X), *car10* (SALK_122977C) T–DNA insertion mutants were obtained from the Salk Institute (http://signal.salk.edu). The double mutants *car56*, *car59* were respectively obtained by crossing *car5* (SAIL_802_B08) with *car6* (SALK_052572.54.00.x) and crossing *car5* (SAIL_802_B08) with *car9* (SALK_088115.56.00.X), then confirmed by PCR with specific primers (Table S1). The *car145*, *car149*, *car459* and *car159* triple mutants were previously described (Rodriguez *et al*., 2014). The *car1459* mutant was generated in this work by crossing *car145* with *car149* and genotyping of the F2 seeds with primers described by Rodriguez et al., (2014) (Rodriguez *et al*., 2014). Next, several *car14569* pentuple mutants were generated in Dr. Pedro L. Rodriguez’s laboratory via CRISPR–Cas9 knockout of the *CAR6* gene in the *car1459* mutant background. Several *car6* alleles were thus obtained. The *car6* allele (termed *car6*–Δ20) in *car14569*–Δ20 loses the sequence “GAAGGAACTGGTAGGGCTTG” (from 30–49 bp) respect to *CAR6*; the *car6* allele (termed *car6*–Δ10) in *car14569–*Δ10 loses the sequence “AACTGGTAGG” (from 36–47 bp), the *car6* allele (termed *car6*–Δ4) in *car14569–*Δ4 loses the sequence “AGGG” (43–46 bp) and the *car6* allele (termed *car6*–Δ1) in *car14569–*Δ*1* loses the nucleotide 43. A single guide RNA (sgRNA) targeting *CAR6* close to the N–terminus was designed using the online tool CRISPR–PLANT (http://www.genome.arizona.edu/crispr/CRISPRsearch.html). Thus, a 19-bp sequence (GAGGTTGAGATGAAGGAAC) followed by the TGG PAM sequence was selected and cloned as a guide-RNA into the entry vector pMTN2982 (GenBank accession number MG917725.1). The sgRNA driven by Arabidopsis U6–26 promoter was cloned into the pHEE2E-TRI vector (Addgene Plasmid #71288), which expresses Cas9 driven by the egg cell-specific EC1.2 promoter (Wang *et al*., 2015). Transformed plants were selected in medium MS supplemented with 20 μg/mL hygromycin and a CAR6 fragment was amplified and sequenced using the following primers: CAR6g-F (AGTAGTAGAGTCAGGAGACAGG) and CAR6g-R (GATTCTTTGGTCTTCCATCTGCG).

For overexpression assays, full–length coding sequences of *CAR4*, *5*, *6*, *9*, *10* respectively fused with a C–terminal Myc tag driven by the ACT2 promoter were cloned into pDT1(Zhu *et al*., 2020). We obtained transgenic plants *CAR1,4,5,6,9,10– Myc* in WT (Col–0) background by *Agrobacterium*–mediated transformation. The transgenic plant CAR9–Myc/*fer–4* was obtained by *Agrobacterium*–mediated transformation of CAR9–Myc in *fer–4* background. In the F2 generation, the *fer–4* background was genotyped by PCR using gene–specific primers (Table S1). Wild–type (Col–0), *fer–4* (Duan *et al*., 2010), and *35S::YFP–CAR9* (Qin *et al*., 2019) were previously described.

### Pull–Down Assay

The full–length *CARs* CDS were cloned into the pGEX–4T–1 and transformed into *E. coli* BL21 to express GST–CAR fusion protein at 16 °C. The GST–tagged CAR proteins were purified as described in the manufacturer’s manual using Pierce Glutathione Agarose (16102, Thermo Fisher Scientific, USA). The 6 × His–tagged FER–CD were purified as described by (Li *et al*., 2018). Recombinant FER–CD protein was incubated overnight at 4 °C with GST beads coupled with GST–CAR in the binding buffer [50 mM Tris–HCl (pH 7.5), 150 mM NaCl, 5 mM MgCl_2_]. The beads were washed three times with the washing buffer [50 mM Tris (pH 7.5), 150 mM NaCl, 0.1 % Triton X–100] and boiled the beads in SDS–PAGE loading buffer, and eluted proteins were analyzed by immunoblot with anti–His (M20001, Abmart) or anti–GST (SC– 80998, CMC) antibody.

### Bimolecular Fluorescence Complementation (BiFC) Assay

The full–length CDS of *FER* and *CARs* were cloned into plasmid pE3308 and pE3449, respectively (Du *et al*., 2016). Protoplasts were isolated from 4–week–old *Arabidopsis* rosette leaves through cellulase and macerozyme digestion for 3 hours. Then, protoplasts were co–transfected with 20 μg FER–nVenus and CAR–cCFP constructs or negative control constructs by the polyethylene glycol transformation method as previously described. The transfected protoplasts were incubated in the dark at 22 °C for 12–16 hours. Fluorescence was monitored with a confocal microscope using an excitation wavelength of 488 nm for GFP and 560 nm for FM4–64 dye (Red).

### *In vitro* Phosphorylation Assay

The phosphorylation and dephosphorylation assays were described as previously (Du, *et al*., 2016). GST–CAR was purified by GST–beads. FER–CD–His and FER^K565R^– CD–His (kinase dead form) were purified by Ni–beads. 1 μg recombinant proteins (GST–CAR and His–FER–CD/His–FER^K565R^–CD) were mixed and added into reaction buffer [50 mM Tris–HCl (pH 7.5), 10 mM MgCl_2_, 1 mM CaCl_2_, 1 mM ATP, 1 mM DTT] to a total volume of 50 μL. The mixture was incubated at 37 °C for 30 min. The dephosphorylation assay was performed by CIP (EF0651, Thermo Fisher Scientific, USA). The reaction was stopped by adding 2 × SDS loading buffer. Proteins were separated on a 10 % SDS–PAGE gel and analyzed by anti–His or anti–GST antibody.

The ABA–induced phosphorylation co–expression system was described previously (Li *et al*., 2018). Vectors of pACYC–PYL1–FER (S–tag), pACYC–PYL1– FER^K565R^ (S–tag), and pRSF–ABI1–CAR9 (His–tag) were constructed. pRSF–ABI1– CAR9 together with pACYC–PYL1–FER (or pACYC–PYL1–FER^K565R^) were co– transformed into one BL21 (DE3) *E. coli* strain. The transformed *E. coli* were cultured at 37 °C until the OD_600_ reached 0.6. Then, 250 μM isopropyl–β–d–thiogalactoside (IPTG) was added to induce the protein expression for 3 hours before 50 µM ABA was added to the bacterial culture to release the FER phosphorylation activity for 10 minutes. After co–expression with FER–CD or FER^K565R^–CD, His–CAR protein was purified and then subjected to alkylation/tryptic digestion followed by liquid chromatography– tandem mass spectrometry (LC–MS/MS). For detail, 10 % SDS–PAGE gel was stained by coomassie G–250. The target bands were isolated and placed into clear tubes and digested by trypsin. Mass spectrometry was performed as described previously. Mass spectrometry was performed using LTQ–Orbitrap. Raw data were analyzed by Xcalibur v.2.1 (Thermo Scientific, Waltham, MA, USA) and Proteome Discoverer v.1.3 beta (Thermo Scientific, Waltham, MA, USA) (*Arabidopsis* database).

### *In vivo* Phosphorylation Assay

Phos–tag assay was performed as described previously (Chen *et al*., 2018). Seven–day– old *CAR–Myc* seedlings (about 0.1 g) were vertically grown on 1/2 MS medium which was solidified by 1.2 % agar. We added more agar in order to avoid the entrance of roots and root hairs to medium (it will hurt plant when separated them from medium). In order to avoid the disturbance of phosphorylation levels that may have been caused during removal from the solid medium. The plants on the medium were soaked by 1 μM RALF1 peptide for 10 min. After that, the plants were quickly moved in liquid nitrogen and grounded to fine powder before mixing with 70 μL 1 × SDS–PAGE loading buffer for 10 min boiling. Centrifuged at 12,000 g for 10 min and transfer the supernatant to a new tube. 10 μL supernatant was saved as input. For the detection of phosphorylated CAR proteins, 20 μL sample was separated by 10 % SDS–PAGE with 35 μM Phos–tag (WAKO, AAL–107) under lower voltage (80 v). PVDF membrane was soaked in methanol for 5 min before use, and then transferred the membrane under 100 mA for 2 h on ice (low temperature is important). The PVDF membrane was blocked with 5 % skim milk for 2 h, and then detected with anti–Myc (CST, 2276S) antibody.

For phosphorylation sites detecting *in vivo*, 2 g of *CAR–Myc* treated with or without 1 μM RALF1 for 10 min as described above. Total protein was extracted by using 0.5 mL of NEB buffer [20 mM Hepes–KOH (pH 7.5), 40 mM KCl, 1 mM EDTA, 1 mM PMSF, 1 % protease inhibitor mixture (Thermo Fisher Scientific, 78420), and 1 % phosphatase inhibitors (Bimake, B15001)]. The homogenized sample was centrifuged at 12,000 g for 10 min each at 4 °C. Then, 20 µL prepared anti–Myc magnetic beads (B26301) for each sample was used to immunoprecipitate CAR–Myc protein complexes at 4 °C for 5 hours. The beads were washed three times in washing buffer [20 mM Hepes–KOH (pH 7.5), 40 mM KCl, and 0.1 % Triton X–100] at 4 °C. The beads were resuspended in 30 µL of 1 × SDS loading buffer, boiled for 10 min, separated by 10 % (w/v) SDS–PAGE gel. The CAR–Myc bands digested and analyzed by MS as described previously.

### Di–4–ANEPPDHQ Lipid Staining

For di–4–ANEPPDHQ lipid staining, four–day–old seedlings are collected for each genotype. 5 μM (work concentration) di–4–ANEPPDHQ (Thermo Fisher, D36802) was added to the liquid 1/2 MS medium for 5 min staining, then washing 3 times using liquid 1/2 MS medium. The seedlings were observed on the Zeiss LSM 880 (Alpha Plan Apochromat 63 ×, NA 1.4 oil objective). Quantification of lipid polarity in root cell meristematic zone has been described previously (Huang et al., 2019; Owen et al., 2011). Briefly, the di–4–ANEPPDHQ fluorescence signal was excited at 488 nm, the membrane order phase was recorded in the range of 500 – 580 nm, the membrane disorder phase was recorded in the range of 620 – 750 nm. The signal of di–4– ANEPPDHQ staining was quantified by ImageJ. For the quantification, both ordered (500 – 580 nm) and disordered (620 – 750 nm) phase fluorescence images are designated as ch00 and ch01, respectively. The GP (Generalized Polarization) value, which is positively correlated with the degree of membrane lipid order, was obtained by calculating mean value of all effective pixels recorded in each region of interest (ROI) of the image. The analysis threshold is fixed at 15, the color scale of the output GP image is set to "16 colors", and no immunofluorescence mask was selected. After processing, manually select the ROI from the GP image for further calculation of the average GP value. More than 40 ROIs were selected from at least 6 images of each process to generate average GP value. The representative images in this work showed a “merge” image and a “HSB” image. “Merge” is merged of the green (500 – 580 nm) and red (620 – 750 nm) emission channels from di–4–ANEPPDHQ. “HSB” (hue– saturation–brightness) picture, a GP image with radiation color coding. The GP value is specified as hue (color) and the average intensity is set as brightness.

### ROS Burst Assay

The flg22*–*induced ROS burst measurement assay was performed as previously described (Stegmann *et al*., 2017). Four–week–old *Arabidopsis* rosette leaves were cut into pieces (4 mm in diameter) and collected in 96–well plates containing sterile water (pH 5.8) and recovered overnight in dark at 22 °C. After 24 hours, sterile water was replaced by a reaction solution [20 μm/L luminol L–012 (Sigma–Aldrich), 1 μg/mL horseradish peroxidase (HRP, Sigma Aldrich), 0.1 μm/L flg22] with or without 1 μM RALF1 (RALF1 elution buffer as a negative control). Luminescence was measured for the indicated time period using Fluoroskan Ascent FL (Thermo Scientific). For the MβCD treatment, we first prepared 10 mM MβCD solution (1.331 g powder dissolved in 100 ml sterile water, pH 5.8) (Solarbio, M9040) and pretreated sample for 30 min. Next, we replaced the MβCD solution with reaction solution and the following steps were as previously described. ROS production was displayed as either the progression of photon counts or the integration of total photon counts. There were three replicates of every sample. Values are means of the total photon counts inhibition over 25 min ± SD. We exported the original data from Fluoroskan Ascent FL and processed data through GraphPad Prism software.

ROS fluorescent probe H2DCF-DA (2′,7′-dichlorodihydro-fluorescein diacetate, Sigma) staining (James *et al*., 2015) 6-day-old seedlings were treated with the liquid 1/2 MS medium containing 10 μM H2DCF-DA for 10 min, the seedlings were washed three times with the culture solution, each for 5 min. Fluorescence was monitored with a confocal microscope using an excitation wavelength of 488 nm.

